# Yin Yang 1 sets up the stage for cerebellar astrocyte maturation

**DOI:** 10.1101/2021.05.14.444129

**Authors:** Karli Mockenhaupt, Katarzyna M. Tyc, Adam McQuiston, Avani Hariprashad, Debolina D. Biswas, Angela S. Gupta, Amy L. Olex, Sandeep K. Singh, Michael R. Waters, Jeff L. Dupree, Mikhail G. Dozmorov, Tomasz Kordula

## Abstract

Diverse subpopulations of astrocytes tile different brain regions to accommodate local requirements of neurons and associated neuronal circuits. Nevertheless, molecular mechanisms governing astrocyte diversity remain mostly unknown. We explored the role of a zinc finger transcription factor Yin Yang 1 (YY1) that is expressed in astrocytes. We found that specific deletion of YY1 from astrocytes causes severe motor deficits in mice, induces Bergmann gliosis, and results in simultaneous loss of GFAP expression in velate and fibrous cerebellar astrocytes. Single cell RNA-seq analysis showed that YY1 exerts specific effects on gene expression in subpopulations of cerebellar astrocytes. We found that although YY1 is dispensable for the initial stages of astrocyte development, it regulates subtype-specific gene expression during astrocyte maturation. Moreover, YY1 is continuously needed to maintain mature astrocytes in the adult cerebellum. Our findings suggest that YY1 plays critical roles regulating cerebellar astrocyte maturation during development and maintaining a mature phenotype of astrocytes in the adult cerebellum.

## Introduction

Astrocytes tile the entire central nervous system and play critical functions by metabolically supporting neurons, maintaining ion balance, regulating concentrations of neurotransmitters, regulating the blood brain barrier, reinforcing and pruning synapses, guiding migrating neurons, and aiding with immune function (Farmer & Murai, 2017, Khakh & Sofroniew, 2015). It is now recognized; however, that astrocytes are extremely diverse across different brain regions and perform specialized functions (Farmer & Murai, 2017, Zhang & Barres, 2010). The regional specialization fine-tunes astrocytes to the local requirements of neurons and associated neuronal circuits (Ben Haim & Rowitch, 2017, Chai, Diaz-Castro et al., 2017, Khakh, 2019, Khakh & Sofroniew, 2015). This astrocyte diversity is initially generated by intrinsic patterning programs (Farmer, Abrahamsson et al., 2016, Farmer & Murai, 2017, Ge, Miyawaki et al., 2012, Hochstim, Deneen et al., 2008), and then astrocyte maturation is promoted by region-specific communication with neurons (Farmer & Murai, 2017, Ge et al., 2012, Hochstim et al., 2008, Huang, Woo et al., 2020, Morel, Chiang et al., 2017). However, the mechanisms regulating astrocyte’s specialized functions that accommodate local needs at different brain regions have only begun to be discovered (Huang et al., 2020). Most of the recent studies focused on differences and similarities of astrocytes between brain regions, but little is known about generation of specific astrocyte subpopulations within the same brain region (Batiuk, Martirosyan et al., 2020, Carter, Bihannic et al., 2018, Chai et al., 2017, Gupta, Collier et al., 2018, John Lin, Yu et al., 2017, Lozzi, Huang et al., 2020, Morel et al., 2017, Zeisel, Hochgerner et al., 2018). In the cerebellum, Bergmann glia (BG) of the molecular layer, velate astrocytes (VA) of the granular cell layer, and fibrous astrocytes (FA) of the white matter are the three major astrocyte subpopulations that derive from common progenitors (Buffo & Rossi, 2013, Farmer et al., 2016, Gomez, 2019). Specialization of differentiated BG requires multiple inputs, including induction of Zinc Finger E-Box Binding Homeobox 2 (*Zeb2*) expression (He, Yu et al., 2018) and signaling by Sonic Hedgehog (SHH) secreted by Purkinje cells (PC) (Farmer et al., 2016, Farmer & Murai, 2017). BG acquire highly specific morphology, express AMPA receptors critical for maintaining synapses with PC, and regulate the fine-tuning of motor skills (Saab, Neumeyer et al., 2012). Other mechanisms diversifying cerebellar astrocytes remain enigmatic.

We explored a zinc finger transcription factor, Yin Yang 1 (YY1), which regulates embryogenesis (Donohoe, Zhang et al., 1999), cell cycle (Petkova, Romanowski et al., 2001, Wang, Wu et al., 2018), X chromosome inactivation (Chen, Shi et al., 2016), oncogenesis (Gordon, Akopyan et al., 2006), and differentiation of multiple cell types (Beagan, Duong et al., 2017, Gregoire, Karra et al., 2013, He, Dupree et al., 2007, Kleiman, Jia et al., 2016, Liu, Schmidt-Supprian et al., 2007). YY1 plays diverse roles in transcriptional regulation by interacting with other regulatory proteins (Bengani, Mendiratta et al., 2013, Cai, Jin et al., 2007, Chiang & Roeder, 1995, Lee, Galvin et al., 1995, Rezai-Zadeh, Zhang et al., 2003, Srinivasan, Pan et al., 2005, Wang et al., 2018), including histone deacetylases and acetyltransferases (Atchison, 2014, Weintraub, Li et al., 2017). YY1 was also recognized as an architectural transcription factor that creates active loops of chromatin (Atchison, 2014, Beagan et al., 2017, Weintraub et al., 2017), which provides a common mechanism for many of its diverse functions. Indeed, YY1 orchestrates chromatin looping during early neural lineage commitment (Beagan et al., 2017, Weintraub et al., 2017), is critical for the development of the neuroepithelium (Dong & Kwan, 2020), and maintains proliferation and survival of neural progenitor cells (Zurkirchen, Varum et al., 2019). Although YY1 is also highly expressed in astrocytes (Karki, Webb et al., 2014, Waters, Gupta et al., 2019), relatively little is known about its astrocyte-specific functions and mechanisms of action. In this study, we found that YY1 is critical to execute and sustain unique and distinct gene expression programs in subpopulations of cerebellar astrocytes.

## Results

### YY1 deletion in GFAP^+^ cells triggers severe motor deficits and distinct effects on subpopulations of cerebellar astrocytes

Although YY1 is ubiquitously expressed in the central nervous system and is critical in neural progenitors (Beagan et al., 2017, Weintraub et al., 2017, Zurkirchen et al., 2019) and oligodendrocytes (He et al., 2007), its biological functions in astrocytes remain poorly understood. To delineate YY1’s roles in astrocytes *in vivo*, we generated conditional knock-out mice with the *Yy1* gene specifically deleted from GFAP^+^ cells (*Yy1^ΔAST^*) using *mGfap-cre* mouse line that drives CRE expression in the GFAP^+^ cells, including astrocytes in the cerebellum starting at P7 (Tao, Wu et al., 2011). As expected, only partial depletion of YY1 was observed in the brain lysates of *Yy1^ΔAST^* mice (Fig. 1A), suggesting a successful *Yy1* deletion in astrocytes, but not other cell types. Accordingly, YY1 was abundant in cultured astrocytes of control littermates but not expressed by astrocytes derived from *Yy1^ΔAST^* mice (Fig. 1B). Specific deletion of YY1 was also evident in cerebellar astrocytes *in vivo* already at postnatal day 10 (Fig. 1C). In contrast to astrocytes, YY1 was expressed by cultured neurons of both control and *Yy1^ΔAST^* littermates (Fig. 1D), emphasizing the specificity of YY1 deletion. At birth, the *Yy1^ΔAST^* mice were indistinguishable from their littermates, which aligns with postnatal development of astrocytes. However, as these mice aged, *Yy1^ΔAST^* animals gained weight more slowly, remained smaller into adulthood (Fig. 1E), and had a drastically shorter lifespan (Fig. 1F). *Yy1^ΔAST^* mice developed severe motor deficits characterized by a slower velocity (Fig. 1G), an increased number of paw slips on a balance beam (Fig. 1H, and Supplementary Videos 1-8), and a widened gait (Fig. 1I).

**Fig. 1.**
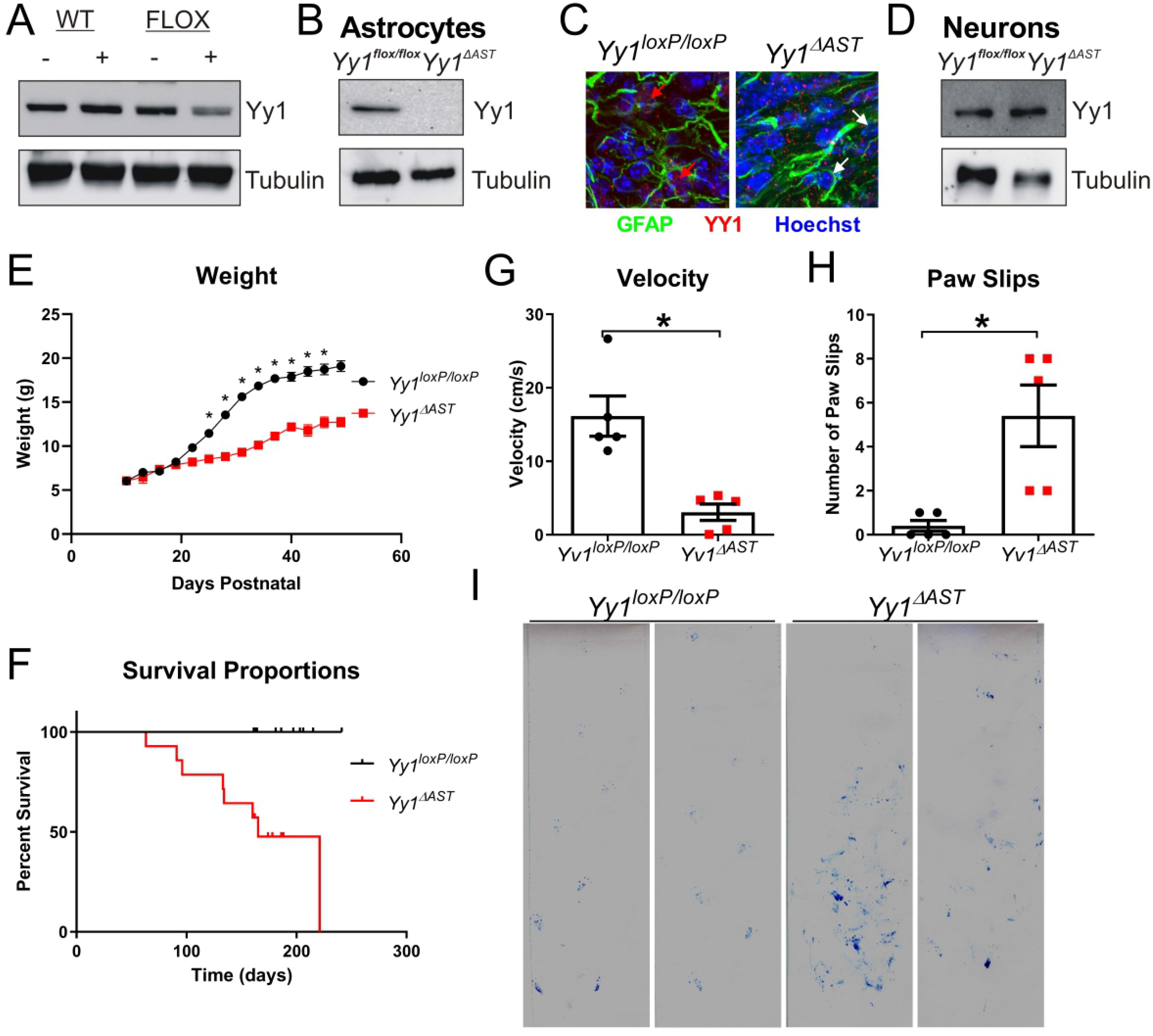
Deletion of YY1 in astrocytes drastically affects motor function. (**A-D**) Expression of YY1 protein in (**A**) total brain lysates, (**B**) cultured astrocytes, (**C**) P10 mouse cerebellum, and (**D**) cultured cerebellar neurons. (**E**) Mean weight of *Yy1^loxP/loxP^* and *Yy1^ΔAST^* mice (n=15, 9; two-tailed t-test; * p <0.0001). (**F**) Survival of *Yy1^loxP/loxP^* and *Yy1^ΔAST^* mice (n=13, 14; Log Rank, Mantel-Cox Test; p=0.005). (**G**) Mean velocity (n=5, 5; two-tailed t-test; p=0.002), and (**H**) mean number of paw slips as mice crossed a balance beam (n=5, 5; two-tailed t-test; * p=0.008). (**I**) Tracks of *Yy1^loxP/loxP^* and *Yy1^ΔAST^* mice crossing narrow path.

Although motor deficits can be associated with disfunction of several regions of the central and peripheral nervous systems, cerebellar neurodegeneration was evident in 5 month-old *Yy1^ΔAST^* mice (Supplementary Fig. 1). The cerebellum regulates muscle synergy by integrating signals from the peripheral nervous system (Leto, Arancillo et al., 2016). Cerebellar insults result in ataxia, a widened gait, attention tremor, PC dysfunction, and learning deficits (Sathyanesan, Kundu et al., 2018). During development, cerebellar cells migrate from the external granule layer to their proper positions, establishing the intricate cytoarchitecture. Initially, PCs migrate along radial glia and form the PC layer (Buffo & Rossi, 2013). Radial glia differentiate into BG relocating their cell bodies into the PC layer, withdrawing from the ependymal layer, retaining their end feet in the pial basement membrane, and enwrapping PC synapses. Subsequently, Granule Cells (GC) migrate across the BG, composing the thick GC Layer (GCL). Radial glia also differentiate into VA of the GCL and FA of the white matter (WM) (Cerrato, 2020). Despite motor deficits evident in adult (8-12 weeks old) mice, we found no substantial differences in brain morphology (Fig. 2A) or weight of *Yy1^loxP/loxP^* and *Yy1^ΔAST^* cerebella (Fig. 2B). There was also no difference in thickness of the ML, GCL, or WM (Supplementary Fig. 2). However, there was a dramatic increase in reactive astrocyte marker expression (Zamanian, Xu et al., 2012), including GFAP, SERPINA3N, and LCN2 as well as a decrease in SPARCL1 expression (Fig. 2C), indicating presence of reactive astrocytes in the *Yy1^ΔAST^* cerebella. In contrast, expression of neuronal (NRG1 and RBFOX3) (Fig. 2D) and oligodendrocyte markers (MBP and PLP1) (Fig. 2E) were not affected, suggesting activation of astrocytes but not a loss of neurons and oligodendrocytes. Further analysis of the cytoarchitecture revealed extensive astrogliosis (Bergmann gliosis) only in the ML of *Yy1^ΔAST^* mice (Fig. 2F) indicated by GFAP^+^ reactive hypertrophic BG exhibiting fewer processes that were disorganized (Fig. 2G). In striking contrast to BG, there was decreased abundance of GFAP^+^ VA and FA in the GCL and WM of *Yy1^ΔAST^* mice, respectively (Fig. 2F). In addition to the distinct effects on astrocytes, calbindin staining revealed reduced arborization of PC (Fig. 2F); however, the number of PC bodies was unchanged (Fig. 2H). Furthermore, the numbers of Iba1-positive cells were significantly increased (Fig. 2F, I), suggesting ongoing inflammation. Taken together, these findings suggest that astrocytic YY1 exerts unique effects on subpopulations of astrocytes that may be critical for the coordination of motor functions.

**Fig. 2.**
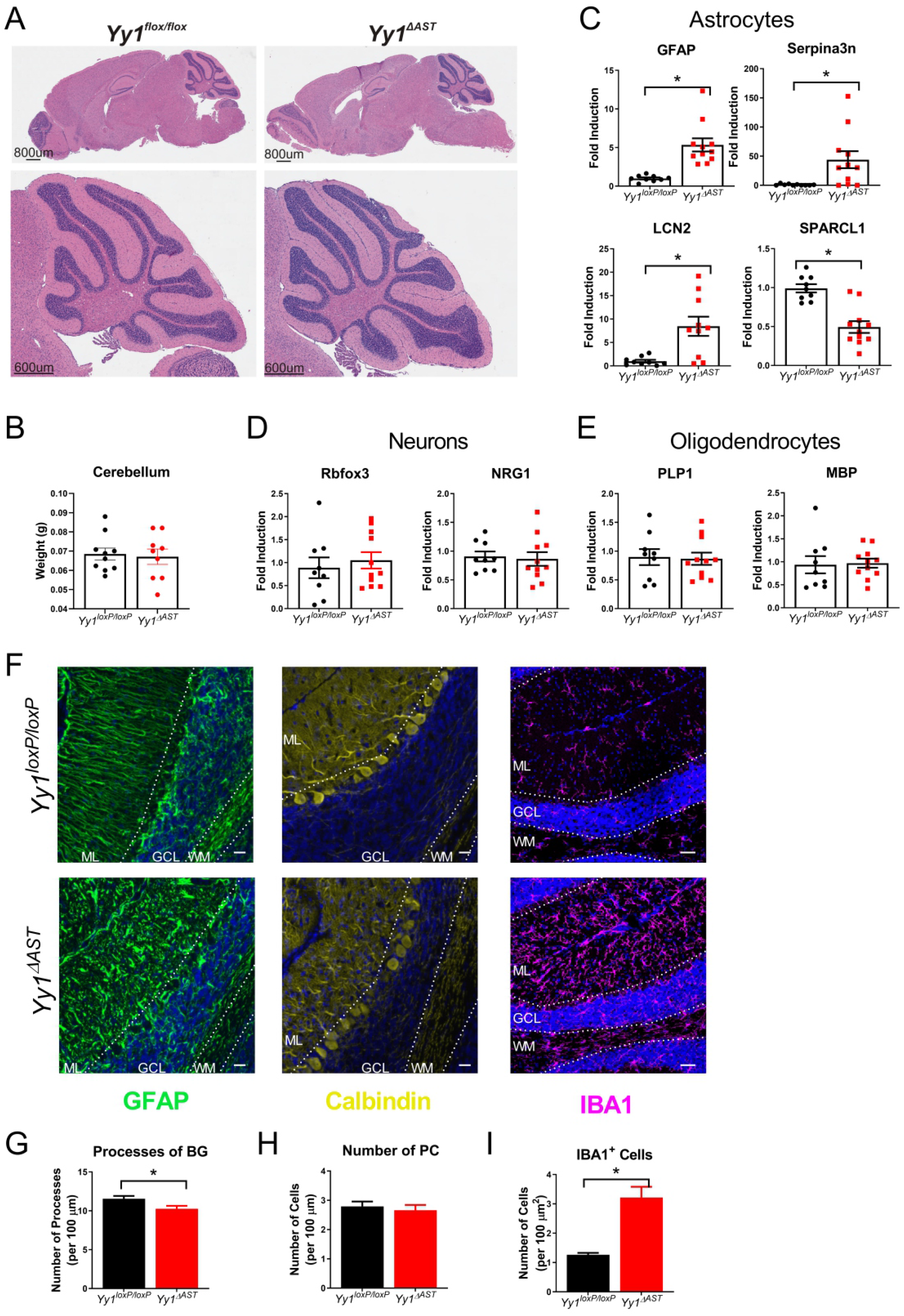
Effect of YY1 deletion on subpopulations of astrocytes in adult cerebellum. (**A**) Hematoxylin and Eosin staining of *Yy1^loxP/loxP^* and *Yy1^ΔAST^* 8-12 week-old brains. (**B**) Weights of *Yy1^loxP/loxP^* and *Yy1^ΔAST^* cerebella (n=10, 9; two-tailed t-test; p=0.78). (**C-E**) Expression of mRNAs (normalized to GAPDH) in cerebella of *Yy1^loxP/loxP^* and *Yy1^ΔAST^* mice (n=9, 11; two-tailed t-test; * p<0.01). (**F**) Immunofluorescence of *Yy1^loxP/loxP^* and *Yy1^ΔAST^* cerebellum with GFAP, Calbindin, Iba1, and Hoechst. Confocal images at 40X, scale bar 20 μm. ML, molecular layer; GCL, granular cell layer; WM, white matter. (**G**) Quantification of GFAP^+^ processes in the ML (n=50, 30 locations; two-tailed t-test; * p=0.02). (**H**) Quantification of PC bodies (n=29, 29 locations; two-tailed t-test; p=0.600). (**I**) Quantification of IBA1^+^ cells (n=35, 18 locations; two-tailed t-test; * p<0.0001).

### YY1 is continuously needed in astrocytes

YY1 could be temporarily needed during astrocyte development or permanently required in mature cerebellar astrocytes. To address the later possibility, we generated tamoxifen (TAM)-inducible astrocyte-specific YY1 cKO mice (*Yy1^Aldh1L1-CreERT2^*) expressing CreERT2 driven by a highly specific *Aldh1L1* promoter (Srinivasan, Lu et al., 2016). We examined the effects of TAM or vehicle administration in adult mice (Fig. 3A). Similarly, to the *Yy1^ΔAST^* mice, *Yy1^Aldh1L1-CreERT2^* mice had dramatically shorter life span after TAM treatment (Fig. 3B). They also displayed motor deficits with a slower velocity (Fig. 3C) and made more mistakes on a balance beam (Fig. 3D). We also found evident Bergmann gliosis in the ML indicated by both thickened processes (Fig. 3E) and decreased numbers of BG processes (Fig. 3F). This was accompanied by a reduction in PC arborization (Fig. 3E). Although the numbers of Iba1^+^ cells have not changed (Fig. 3G), the ameboid morphology of these cells (Fig. 3E, H) indicated their activation. As for the cerebellar VA and FA, no significant changes were detected 18 days after induction. However, analysis of these subpopulations was not possible at later time points because of the short life span of the *Yy1^Aldh1L1-CreERT2^* mice after tamoxifen treatment. These data suggest that continuous expression of YY1 in BG and likely other subpopulations of astrocytes is required to maintain astrocyte specialization even in adult mice.

**Fig. 3.**
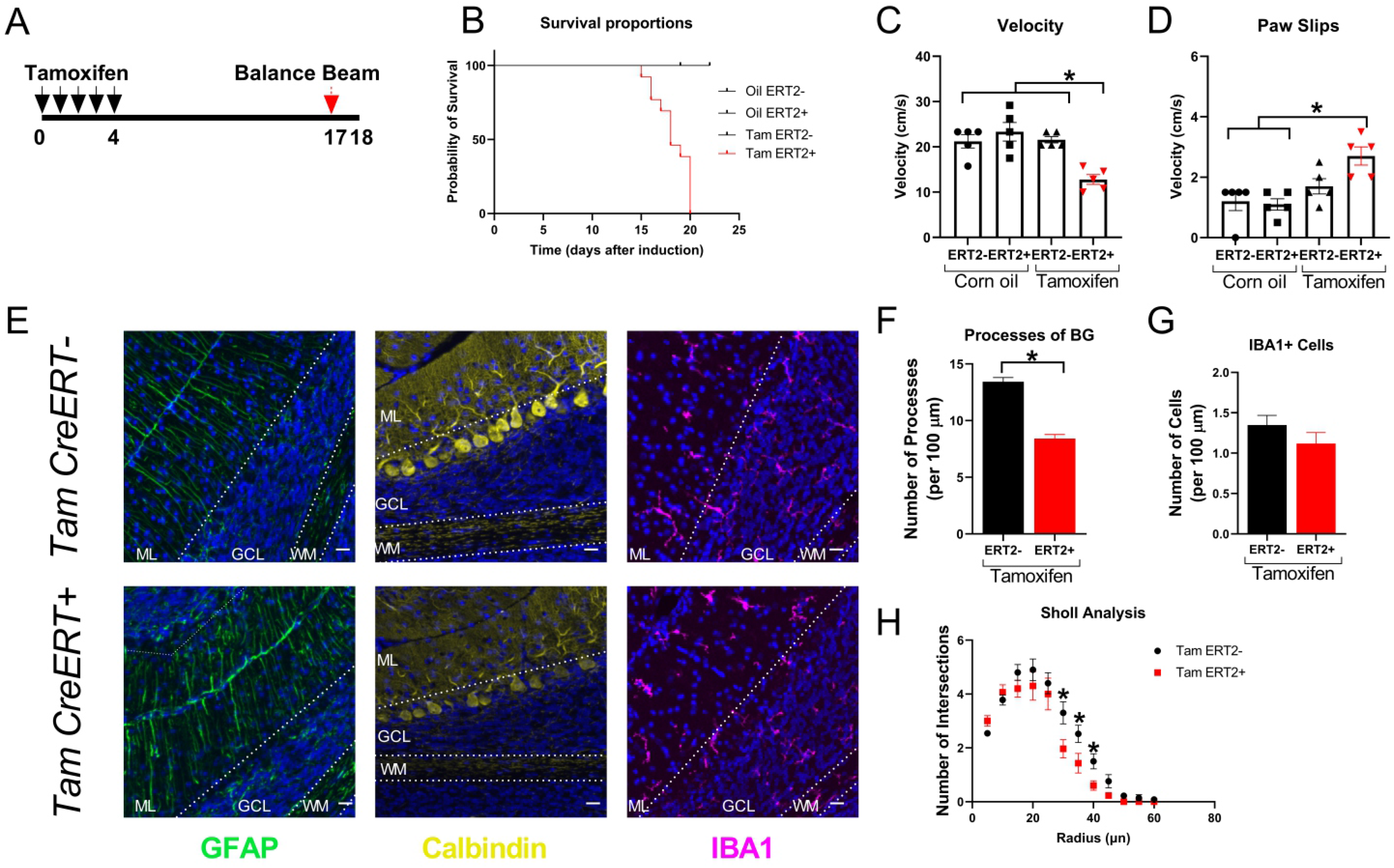
Deletion of YY1 in astrocytes of adult mice. (**A**) Model of experimental design. (**B**) Survival (n=5, 10, 8, 13 mice; Log Rank, Mantel-Cox test; * p<0.0001). (**C**) Mean velocity (n=5, 5, 5, 7; Two-way Anova; * p<0.003). (**D**) Mean number of paw slips on a balance beam (n=5, 5, 5, 7; Two-way Anova; * p<0.003). (**E**) Immunofluorescence of cerebellum (day 18 post induction) stained with GFAP, Calbindin, IBA1, and Hoechst; 20X. ML, molecular layer; GCL, granular cell layer; WM, white matter. (**F**) Quantification of GFAP^+^ processes (n=42, 54 locations; two-tailed t-test; * p<0.0001). (**G**) Quantification of IBA1^+^ cells (n=51, 30 locations; two-tailed t-test; p=0.23). (**H**) Sholl analysis of IBA1^+^ cells (n=51, 30 locations; multiple unpaired two-tailed t-test; * p<=0.03).

### YY1 is dispensable for the initial stages of astrocyte development

Next, we wondered at what timepoint is YY1 required for astrocyte development. First, YY1 could be needed to execute the initial intrinsic patterning programs affecting astrocyte progenitors (Hochstim et al., 2008). In this case, one would expect to find effects of YY1 deletion at early stages of cerebellar development that would manifest in improper cell migration, changes in morphology of immature astrocytes, and changes in gene expression profiles. Alternatively, YY1 could affect subsequent maturation locally promoted by region-specific crosstalk of immature astrocytes with neurons supporting neural circuit assembly (Chai et al., 2017, Farmer et al., 2016). In this scenario, YY1 would be required for latter stages of astrocyte development, whereas the initial steps would be unaltered. We first analyzed cytoarchitecture of cerebella of *Yy1^Aldh1L1-CreERT2^* mice after TAM or vehicle administration at early stages of postnatal development (Fig. 4A). In contrast to the findings in adult *Yy1^Aldh1L1-CreERT2^* mice, there were no substantial changes in the distribution or appearance of astrocytes, PC, or microglia (Fig. 4B). The numbers of BG processes were unaffected (Fig. 4C). Furthermore, the numbers of Iba1^+^ cells were comparable (Fig. 4D), suggesting that YY1 is not needed at early stages of astrocyte development. To verify these findings, we examined cytoarchitecture of *Yy1^loxP/loxP^* and *Yy1^ΔAST^* cerebella at P10. In contrast to what we found in the adult *Yy1^ΔAST^* cerebellum (Fig. 2), astrogliosis was also not evident at P10 (Fig. 4E), and there was no difference in the distribution or appearance of astrocytes, PC, or microglia (Fig. 4E-H). Moreover, the electrophysiological properties of PC were not changed in *Yy1^ΔAST^* mice at P10 (Fig. 4I, J). Quantitative measurement of the action and membrane potential showed no statistical difference in resting membrane potential, input resistance, depolarizing sag to hyperpolarizing current injection, action potential amplitude, duration, or threshold between PC from *Yy1^loxP/loxP^* and *Yy1^ΔAST^* cerebella (Supplementary Table 1). Altogether, the morphology of immature astrocytes and cerebellar cytoarchitecture were initially unaltered in both *Yy1^Aldh1L1-CreERT2^* and *Yy1^ΔAST^* mice. Given that YY1 deletion had no apparent effects on cerebellar cytoarchitecture at P10, we set on to evaluate whether deletion of YY1 from astrocytes manifests in initial changes on the molecular level. To this end, we examined gene expression using single-cell RNA sequencing (scRNA-seq) of *Yy1^ΔAST^* cerebella. In total, we sequenced 11,226 cells and applied unsupervised clustering that identified 16 unique clusters of cells that were shared by *Yy1^loxP/loxP^* and *Yy1^ΔAST^* cerebella (Fig. 5A, Supplementary Table 2). All clusters had similar UMI and gene counts except for cluster 9 (Fig. 5B); however, low values for cluster 9 were found in both *Yy1^loxP/loxP^* and *Yy1^ΔAST^* cerebella. To identify astrocytes, we took a two-pronged approach. First, an unsupervised analysis of the top 10 biomarkers for each cluster identified cluster 6 as astrocytes (Supplementary Fig. 3). Second, we performed a supervised analysis of the expression of known astrocyte markers, which indicated that cluster 6 indeed represented astrocytes (Fig. 5C). Furthermore, comparison of gene expression profiles of cluster 6 to previously published profiles of astrocytes derived from different brain regions (Batiuk et al., 2020, Borggrewe, Grit et al., 2021, Zhang, Sloan et al., 2016) indicated substantial similarities (Fig 5D). Nevertheless, cells in cluster 6 were also unique likely because of their immature astrocyte phenotype. In addition to cluster 6, cells in cluster 11 expressed some of the astrocytic markers (Batiuk et al., 2020); however, *Gfap, Aldh1L1,* and many additional astrocyte markers were expressed at very low levels, and thus these cells were not further analyzed. Importantly, *Yy1* expression was specifically lost from astrocytes (cluster 6), but not other cell types in *Yy1^ΔAST^* cerebella (Fig. 5E, F), confirming highly specific deletion of YY1. Three cerebellar astrocyte subpopulations have been previously described (Gupta et al., 2018, John Lin et al., 2017). In line with these findings, unsupervised re-clustering of cells in cluster 6 yielded three major sub-populations of astrocytes (6a, 6b, and 6c) (Fig. 5G, Supplementary Table 2). Biomarker expression analysis of the sub-clusters (Supplementary Table 2) in combination with Allen Brain Atlas *in situ* RNA expression data (https://developingmouse.brain-map.org/) identified sub-cluster 6a as BG, 6b as VA, and 6c as FA (Fig. 5H-K, Supplementary Fig. 4). Interestingly, although *Gdf10* and *Gpr37l1* are known markers of BG (Carter et al., 2018, Gupta et al., 2018, He et al., 2018, Marazziti, Di Pietro et al., 2013), our analysis uncovered *Cyp26b1* and *Hsd11b1* as novel markers of VA, and *Myoc* and *C4b* as novel markers of FA (Fig. 5H-J), which were still uniquely localized at P28 (Supplementary Fig. 4). Significantly, gene expression profiles of the three astrocyte sub-populations were largely unaffected by YY1 deletion (Fig. 5K). YY1 deletion did not affect expression of astrocyte progenitor markers (Zhang et al., 2016) (Supplementary Fig. 5) or mature astrocyte markers (Zhang et al., 2016) (Supplementary Fig. 6) at P10. Furthermore, expression of reactive astrocyte markers (Escartin, Galea et al., 2021) was also not changed (Supplementary Fig. 7), which coincided with the lack of expression of inflammatory mediators by either astrocytes or microglia (Supplementary Fig. 8). Altogether, our analysis of *Yy1^Aldh1L1-CreERT2^* and *Yy1^ΔAST^* cerebella at P10 indicated that YY1 is dispensable for the initial steps of cerebellar astrocyte development.

**Fig. 4.**
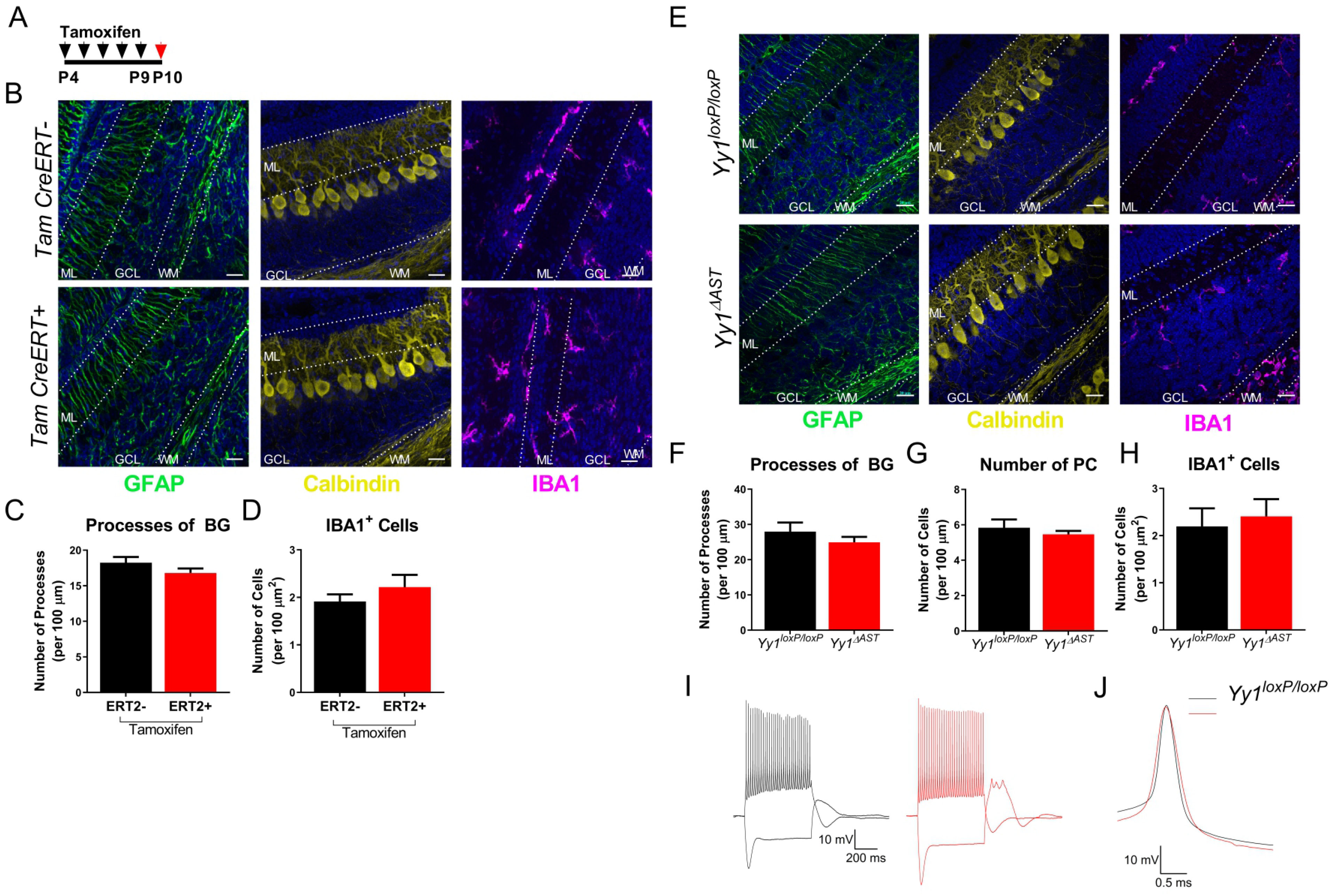
Normal architecture of *Yy1^Aldh1L1-CreERT2^* and *Yy1^ΔAST^* cerebella at P10. (**A**) Model of experimental design. (**B**) Immunofluorescence of TAM-treated cerebella stained with GFAP, Calbindin, Iba1, and Hoechst at 40X, scale bar 20 μm. ML, molecular layer; GCL, granular cell layer; WM, white matter. (C) Quantification of GFAP^+^ processes of BG (n=21, 14; locations; two-tailed t-test; p=0.21). (**C**) Quantification of IBA1^+^ cells (n=5, 5 locations; two-tailed t-test; p=0.34). (**E**) Immunofluorescence of *Yy1^loxP/loxP^* and *Yy1^ΔAST^* cerebellum at P10 stained with GFAP, Calbindin, Iba1, and Hoechst. Confocal images at 40X, scale bar 20 μm. ML, molecular layer; GCL, granular cell layer; WM, white matter. (**F**) Quantification of GFAP^+^ processes of BG (n=21, 26; locations; two-tailed t-test; p=0.29). (**G**) Quantification of PC bodies (n=7, 9 locations, two-tailed t-test; p=0.43). (**H**) Quantification of IBA1^+^ cells (n=33, 38 locations; two-tailed t-test; p=0.69). (**I**) Representative responses of a P10 *Yy1^loxP/loxP^* and *Yy1^ΔAST^* PC to 350 pA depolarizing and -400 pA hyperpolarizing step current injections (600 ms duration). (**J**) Overlay of the first action potentials in the train of E.

**Fig. 5.**
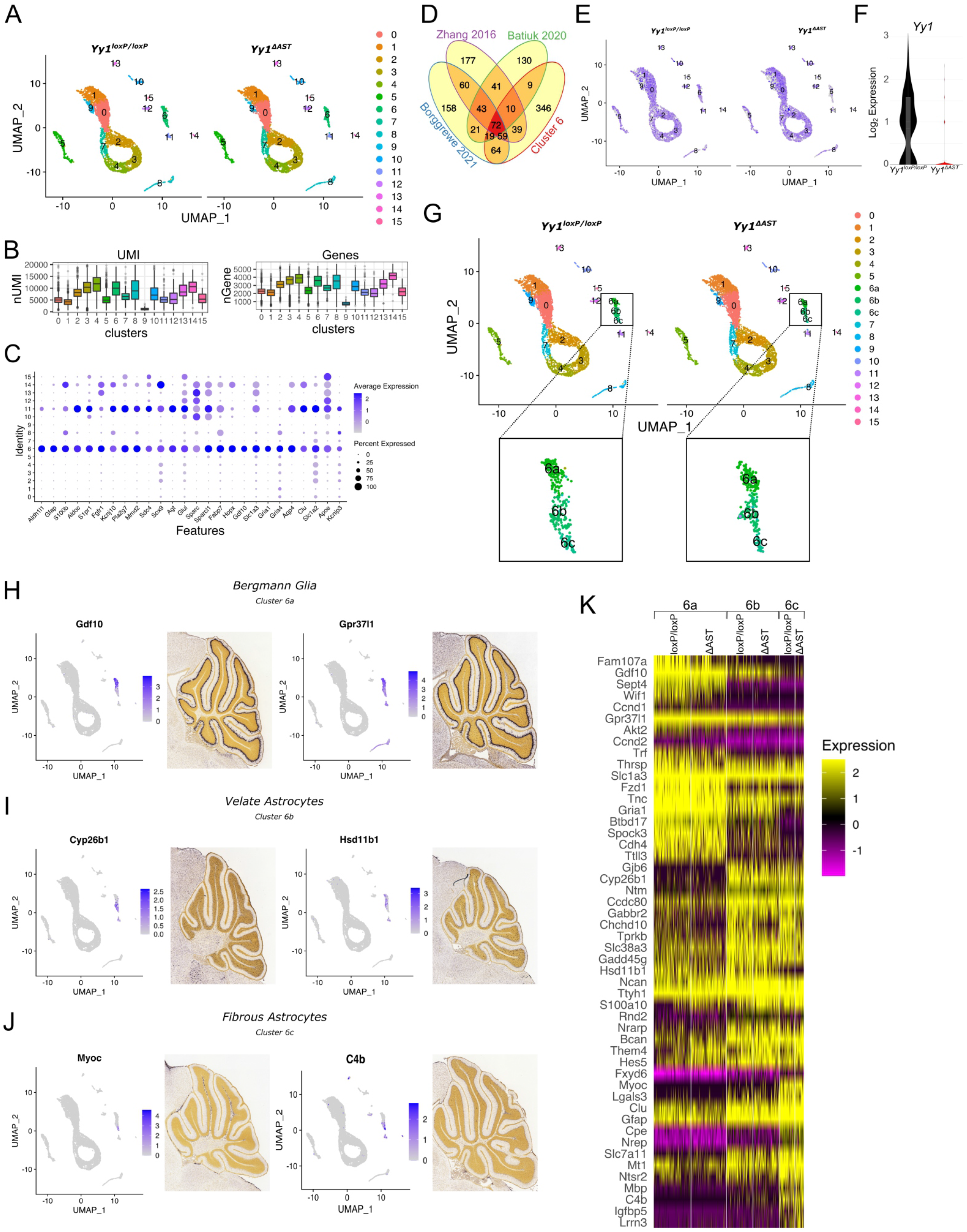
scRNA-seq analysis of *Yy1^loxP/loxP^* and *Yy1^ΔAST^* cerebella at P10. (**A**) Two-dimensional UMAP clustering. 11,226 cerebellar cells (n= 2, 2; mice). (**B**) Box plot of UMI and gene number per cell in each cell cluster. (**C**) Expression of astrocyte markers in each cluster. (**D**) Overlap of top 618 cluster 6 genes with published mouse astrocyte gene sets (Batiuk et al., 2020; Borggrewe et al., 2021; Zhang et al., 2016). (**D**) UMAP and (**E**) violin plot visualization of YY1 expression in *Yy1^loxP/loxP^* and *Yy1^ΔAST^* cerebella. (**G**) Two-dimensional UMAP clustering visualizing subclusters of astrocytes. (**H-J**) UMAP graphs presenting the expression of subpopulation markers and their expression *in situ* (Allen Brain Atlas, P14) in BG (**H**), VA (**I**), and FA (**J)**. (**K**) Heatmap of top 10 biomarkers for each subpopulation. Unsupervised analysis of gene expression profiles of BG, VA, and FA in *Yy1^LoxP/LoxP^* and *Yy1^ΔAST^* cerebella.

### YY1 is required for late stages of astrocyte development

To assess whether YY1 affects subsequent steps of astrocyte development that are governed by region-specific astrocyte-neuron cross-communication (Chai et al., 2017, Farmer et al., 2016), we examined cerebella at P20. In contrast to P10, activation of BG was evident in the ML at P20 in *Yy1^ΔAST^* cerebella while numbers of GFAP^+^ VA and FA were dramatically diminished (Fig. 6A). While processes of BG were affected (Fig. 6B), number of PC bodies remained the same (Fig. 6C). We noticed a decrease in arborization of PC (Fig. 6A); however, electrophysiological properties of PC were not affected (Supplementary Fig. 9, Supplementary Table 3). Activation of BG was also associated with increased amounts of reactive microglia (Fig. 6A and D), indicating an inflammatory environment. Importantly cytoarchitectures of *Yy1^ΔAST^* cerebella at P20 and adult mice (Fig. 2F) were similar, indicating that YY1 exerts its effects on astrocytes at late stages of their development. Both granule cell migration and synapse elimination are completed by P20 (Araujo, Carpi-Santos et al., 2019), and immature astrocytes undergo their location-specific maturation that is driven by signals from the surrounding cells (Araujo et al., 2019).

**Fig. 6.**
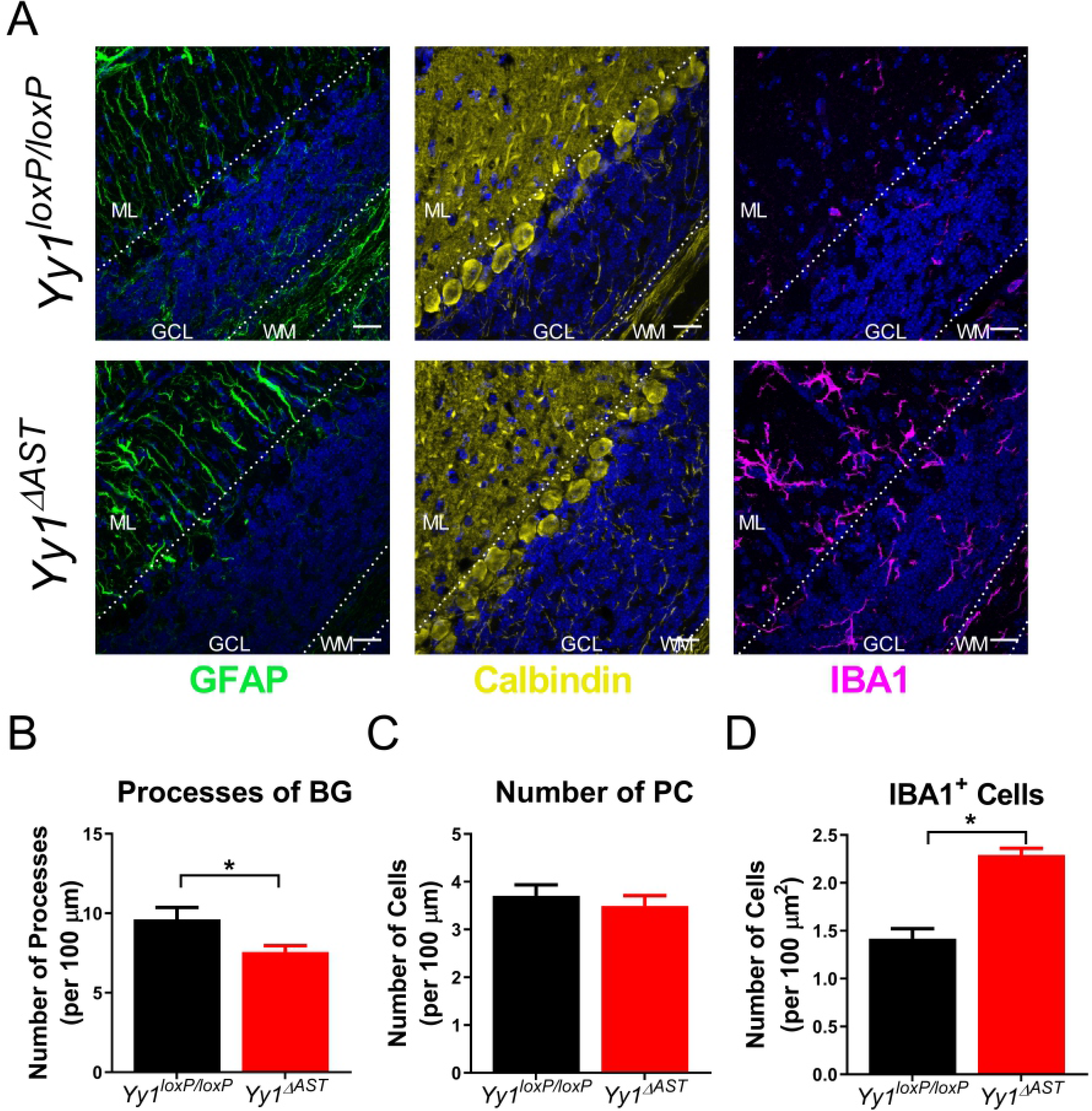
Differential effect of YY1 deletion on subpopulations of cerebellar astrocytes at P20. **(A)** Immunofluorescence of *Yy1^loxP/loxP^* and *Yy1^ΔAST^* cerebella at P20 stained with GFAP, Calbindin, IBA1, and Hoechst. Confocal images at 40X, scale bar 20 μm. ML, molecular layer; GCL, granular cell layer; WM, white matter. **(B)** Quantification of GFAP^+^ processes of BG (n=16, 28 locations; two-tailed t-test; * p=0.01). **(C)** Quantification of PC bodies (n=8, 14 locations, two-tailed t-test; p=0.53). **(D)** Quantification of IBA1^+^ cells (n=19, 25 locations; two-tailed t-test; * p<0.0001).

Subsequently, we established that the initial morphological signs of astrocyte dysfunction occur at P17 (Supplementary Fig. 10A). Since microglia activation was not yet evident at this stage, we reasoned that changes to gene expression between *Yy1^loxP/loxP^* and the *Yy1^ΔAST^* mice at P17 should be attributed to diseased astrocytes and not yet a result of developing inflammation. To analyze gene expression, we sequenced 7,321 cells, applied unsupervised clustering, and identified 25 distinct cell clusters (Fig. 7A, Supplementary Table 4). There were similar UMI and gene counts across all the clusters (Fig. 7B). In stark contrast to what we found at P10, several clusters were unique to either the *Yy1^loxP/loxP^* or the *Yy1^ΔAST^* cerebella. We applied the same two-pronged approach to identify astrocytes. Both the unsupervised analysis (Supplementary Fig. 11) and the supervised analysis (Fig. 7C) identified clusters 8, 10, and 15 as astrocytes in the *Yy1^loxP/loxP^* cerebella, whereas clusters 1 and 9 are astrocytes in the *Yy1^ΔAST^* cerebella. This was in striking contrast to all other cell types that clustered similarly in both the *Yy1^loxP/loxP^* and the *Yy1^ΔAST^* cerebella. Similarly to cluster 11 at P10, cells in cluster 14 at P17 expressed low levels of *Gfap*, *Aldh1L1*, and several other astrocytes markers, and thus were not further analyzed. Importantly, *Yy1* expression was specifically diminished in astrocytes of the *Yy1^ΔAST^* cerebella (Supplementary Fig. 10B, C). Re-clustering of the astrocyte clusters 9 and 10 indicated three major sub-populations present in both clusters, 9a, 9b, and 9c in *Yy1^ΔAST^* cerebella and 10a, 10b, and 10c in *Yy1^loxP/loxP^* cerebella (Fig. 7D and Supplementary Fig. 12). Subsequent analysis of key biomarker gene expression (Fig. 7E-G) indicated that, although unique, both clusters 8 and 15 account for BG, sub-clusters 10a and 10b represent VA, whereas sub-cluster 10c represents FA in *Yy1^loxP/loxP^* cerebella. Similarly to astrocytes at P10, immature astrocyte subpopulations at P17 had unique expression profiles that displayed similarities to previously published profiles of astrocytes (Batiuk et al., 2020, Borggrewe et al., 2021, Zhang et al., 2016) (Fig 7H-J). Furthermore, our analysis of *Yy1 ^ΔAST^* cerebella, identified cluster 1 as BG, sub-clusters 9a and 9b as VA, and sub-cluster 9c as FA. Importantly, while YY1 deletion did not affect astrocyte progenitor marker expression (Zhang et al., 2016) (Supplementary Fig. 13), it dramatically reduced expression of mature astrocyte markers (Zhang et al., 2016) (Supplementary Fig. 14) suggesting that YY1 is needed during astrocyte maturation. This was further reinforced by hierarchical clustering analysis (Supplementary Fig. 10D), which indicated that YY1-deficient astrocytes at P17 are more similar to *Yy1^loxP/loxP^* immature astrocytes at P10 than to *Yy1^loxP/loxP^* astrocytes at P17. Interestingly, expression of reactive astrocyte markers (Escartin et al., 2021) was not suggestive of reactive astrogliosis in the YY1-deficient astrocytes (Supplementary Fig. 15), and there was no induction of expression of inflammatory mediators by astrocytes or microglia (Supplementary Fig. 16), indicating that astrocytes were not yet reactive at P17. However, *Gfap* expression was induced in BG and reduced in both VA and FA of Yy1^ΔAST^ mice (Supplementary Fig. 15). Thus, the loss of GFAP^+^ VA and FA in Yy1^ΔAST^ cerebella (Fig. 2F, 6A) is a result of downregulation of *Gfap* expression and not a decrease of VA and FA. To determine biological effects of YY1 deletion in subpopulations of astrocytes, we performed pathway analysis. Although there were subpopulation-specific effects, in general, processes previously reported to be controlled by YY1 in neural progenitors (Zurkirchen et al., 2019) were increased (Fig. 7K-M), such as cytoplasmic translation” and “ribonucleoprotein complex biogenesis”, and “mRNA metabolic processes”. However, YY1 also reduced many other processes, including those involved in neuronal development, such as “synapse organization” and “regulation of membrane potential”.

**Fig. 7.**
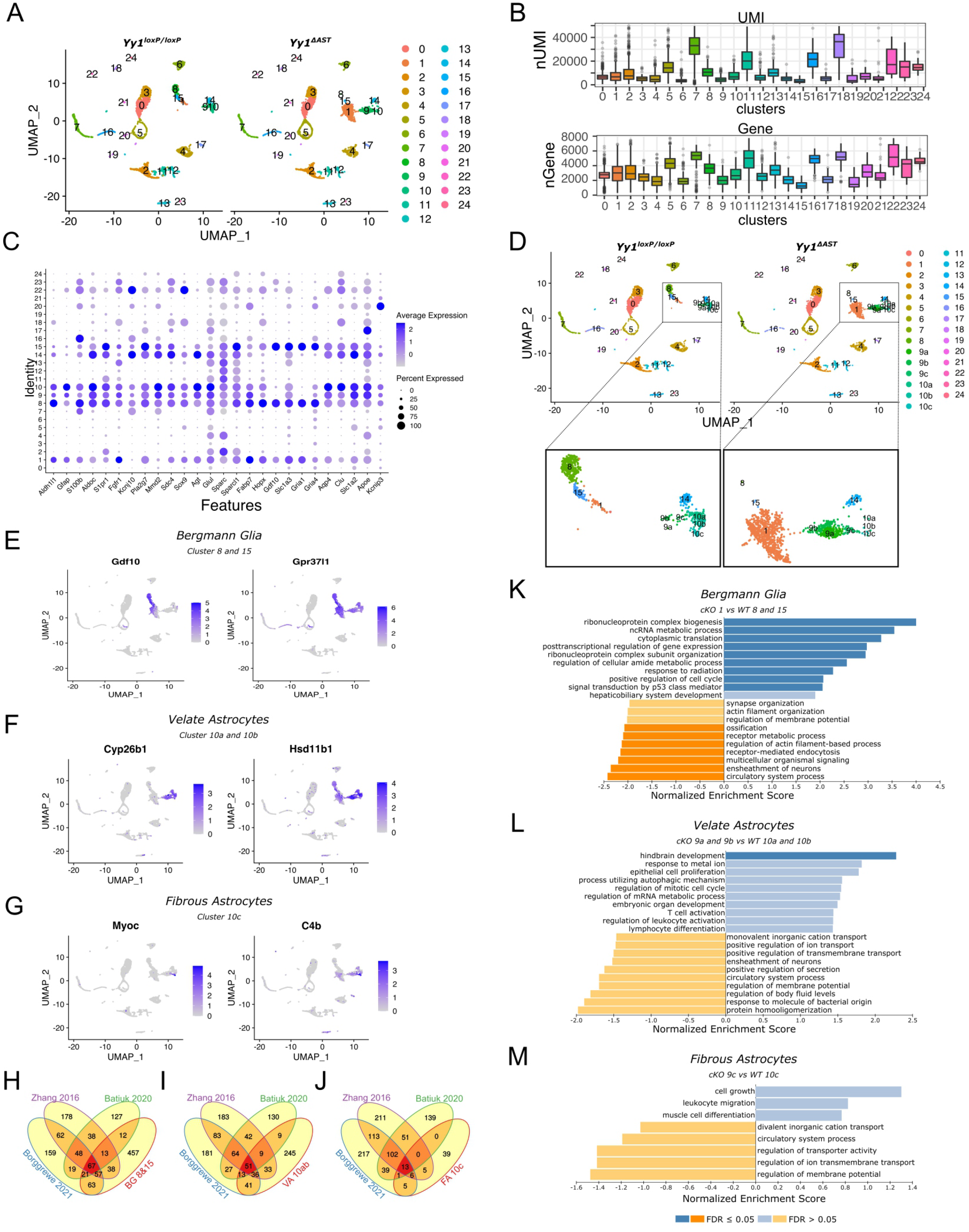
Unique astrocyte clusters are present in *Yy1^ΔAST^* cerebellum at P17. (**A**) Two-dimensional UMAP clustering. 7,321 cerebellar cells were sequenced and analyzed. (n= 2, 2; mice). (**B**) Box plot of UMI and gene number per cell in each cluster. (**C**) Expression of astrocyte markers in each cell cluster. (**D**) Two-dimensional UMAP clustering visualizing subclusters of astrocytes. (**E-G**) UMAP graphs presenting the expression of subpopulation in BG (**E**), VA (**F**), and FA (**G)**. (**H-J**) Overlap of top 728 BG (**H**), 387 VA (**I**), and 208 FA (**J)** genes with published mouse astrocyte gene sets (Batiuk et al., 2020; Borggrewe et al., 2021; Zhang et al., 2016). (**K-M**) Pathway analysis of biological processes regulated by differentially expressed genes in the *Yy1^loxP/loxP^* and *Yy1^ΔAST^* BG (**K**), VA (**L**), and FA (**M**). Top 10 enriched categories are shown.

Together, our data suggest that although YY1 is dispensable during early stages of astrocyte development, it is required during late stages of their development that are governed by region specific interactions with neurons.

## Discussion

Although morphological diversity of astrocytes and their specialized functions have long been recognized (Farmer & Murai, 2017, Meyer, Paisley et al., 2017, Zhang & Barres, 2010), only recently has their molecular diversity been shown using RNA-seq, scRNA-seq, scISOr-seq, slide-seq, and TRAP (Batiuk et al., 2020, Carter et al., 2018, Gupta et al., 2018, John Lin et al., 2017, Rodriques, Stickels et al., 2019). It has been proposed that five molecularly distinct subpopulations of astrocytes occupy specific locations across different brain regions, with three subpopulations representing mature astrocytes (Batiuk et al., 2020). Three major molecularly distinct subpopulations of astrocytes were also found in the cerebellum; however, their identities and precise localizations have not been established (Gupta et al., 2018, John Lin et al., 2017). In this study, we found three major molecular subpopulations of cerebellar astrocytes but, in addition, identified unique markers that allow identification of these subpopulations. Indeed, our analysis identified BG, which were marked by previously known markers *Gdf10*, *Gpr37l1, Slc1a3*, and *Gria1* (Carter et al., 2018, Gupta et al., 2018, He et al., 2018, Marazziti et al., 2013, Saab et al., 2012). Although there was a single cluster of immature BG at P10, we found two distinct molecular subsets of cells that were BG at P17. It is unlikely that these subsets represent BG at different stages of differentiation since neither more closely clusters with immature BG at P10. Importantly, two molecularly unique subpopulations of BG have also been reported by others (Carter et al., 2018). It is possible that one of these BG subsets represents elusive Fañanas cells (Goertzen & Veh, 2018); however, this requires further analysis. In contrast to specific markers of BG, genes specifically expressed in either VA or FA have been difficult to identify to date. Both *Aqp4* and *Bcan* have been used as a marker of VA (Farmer et al., 2016, Yamada, Fredette et al., 1997). While we found that both *Aqp4* and *Bcan* were expressed in VA, *Aqp4* was also upregulated in the FA, and *Bcan* was also expressed in BG, FA, and oligodendrocytes. Our data indicate that in contrast to *Aqp4* and *Bcan*, both *Cyp26b1* and *Hsd11b1* selectively label cerebellar VA at both P10 and P17. However, expression profiles of cerebellar VA do not resemble a specific subpopulation profile of cortical and hippocampal astrocytes (Batiuk et al., 2020), and it remains to be established whether their markers are specifically expressed in other brain regions. Surprisingly, specific markers of FA have not been identified to date. Here we identified *Myoc* and *C4b* to be uniquely expressed in FA but not BG or VA in the cerebellum. Whether these are markers of FA across different brain regions or unique to the cerebellum needs to be determined. These new markers may allow the creation of tools to study subpopulations of cerebellar astrocytes.

While our analysis at P10 indicates that expression profiles of immature BG, VA, and FA are initially different, the profiles of VA and FA were remarkably similar at P17, and these cells clustered together, away from BG. It has been shown that all three cerebellar astrocyte subpopulations stem from common progenitors in the developing cerebellum. Nevertheless, VA and BG are more likely to stem from the same clones (Cerrato, Parmigiani et al., 2018), while FA may derive from their own clones or share a clonal lineage with interneurons (Parmigiani, Leto et al., 2015). Since immature FA and VA are initially different and subsequently their expression profiles become more similar, our findings point to the plasticity of developing astrocytes, and the importance of region-specific external signals that drive maturation of these cells.

Here we have shown that YY1 is needed to execute unique gene expression programs in subpopulations of developing cerebellar astrocytes during their maturation. This is supported by our findings that YY1-deficient astrocytes at P17 fail to gain expression of known mature astrocyte markers and group together with immature astrocytes at P10, indicating a defect in their maturation. YY1 is also permanently required to sustain mature BG and likely VA and FA in adult mice. Thus, YY1 regulates region-specific maturation of cerebellar astrocytes during both development and in the adult cerebellum. Similar findings have recently been reported for another critical transcription factor, NFIA, which also controls region-specific gene expression and maturation of astrocytes, as well as local circuit activities (Huang et al., 2020). These data suggest that both region-specific external signals and intrinsic transcriptional programs are needed to specifically fine tune maturing astrocytes to local needs of the neural circuits. Specific mechanisms by which YY1 exerts its effects in subpopulations of astrocytes remain unknown. Multiple mechanisms of action have been proposed for YY1, including both epigenetic changes to the chromatin and generation of cell-type-specific active chromatin loops (Beagan et al., 2017, Weintraub et al., 2017). Although the chromatin loop model is an attractive mechanism that functions in neural progenitors (Beagan et al., 2017), it remains to be established if YY1 exerts its effects on astrocytes via this or different more traditional mechanisms for a transcription factor. Reorganization of chromatin regulates neural stem cell fate determination, neural plasticity, and learning and memory (Ravi & Kannan, 2013, Sultan & Day, 2011). In contrast, relatively little is known about functions of such reorganization in astrocytes (Galloway, Adeluyi et al., 2018, Ito & Takizawa, 2018) with studies mostly focused on DNA methylation and histone modifications (Neal & Richardson, 2018, Pavlou, Grandbarbe et al., 2019). Our data suggest that both development and maintenance of mature BG, VA, and FA in the cerebellum may be differentially regulated on the chromatin level. This mode of regulation would parallel findings demonstrating distinct chromatin accessibilities in subgroups of neurons (Mo, Mukamel et al., 2015), and further epigenetic alternations associated with neuronal activity (Su, Shin et al., 2017).

Since YY1 has been assumed to date to be dispensable for astrocytes (He et al., 2007), our findings drastically change the understanding of its importance in astrocytes. In summary, our data suggest that YY1 plays a critical role regulating cerebellar astrocyte maturation during the development and maintenance of a mature astrocyte phenotype in the adult cerebellum.

## Acknowledgments

This work was supported by NIH grants R01NS122986, R21NS102802, and R21NS118359 (to TK). Microscopy and histology were performed at the VCU Microscopy Facility and VCU Cancer Mouse Models Core, respectively, supported, in part, by funding from NIH-NCI Cancer Center Support Grant P30 CA016059. This work was also supported by the CTSA award No. UL1TR002649 from the National Center for Advancing Translational Sciences. We would like to thank Mr. Lukasz Kordula for help constructing the balance beam.

## Author Contributions

K.M planned and performed most experiments, with assistance from A.M, A.H., D.D.B., A.S.G., S.K.S., M.R.W., and J.L.D. Bioinformatic analysis was performed by K.M.T. and A.O.L, with supervision of M.G.D. T.K. conceived the study and contributed to planning of the experiments. T.K. and K.M. drafted the manuscript. All authors read, edited, and approved the final manuscript.

## Data Availability

Raw sequencing data are publicly available through the GEO database (GSE166792).

## Materials and methods

### Mice

Mice with the *Yy1* allele flanked by loxP sites (Jackson Laboratory; B6;129S4-*Yy1^tm2Yshi^*/J) were bred with GFAP-Cre (Jackson Laboratory; 77.6mGFAPcre) to generate *Yy1^ΔAST^* mice. *Yy1^loxP/loxP^* mice were also crossed with Aldh1l1-CreERT2 mice (received from Dr. Cagla Eroglu, Duke University) to generate tamoxifen-inducible conditional knockout mice. Mice were housed at Virginia Commonwealth University according to the Institutional Animal Care and Use Committee guidelines. Animals were housed with a 12-hour light/dark cycle, in an animal facility, with standard laboratory chow, and water ad libitum. *Yy1^ΔAST^* mice were given DietGel (ClearH20) once they developed severe symptoms. Randomly chosen littermates (males and females) were used for all experiments, and the group sizes are provided in the figure legends. Survival curves were created from mice that reached a set experimental endpoint (20% weight loss) or died.

### Tamoxifen-induced conditional deletion of Yy1

Tamoxifen was dissolved in 100% ethanol warmed to 55°C (0.2 g/ml). After the tamoxifen was dissolved 9 ml of corn oil (Sigma) was added, incubated at 37°C for 15 minutes, and protected from light. The dissolved tamoxifen was stored at -20°C. Next, 75 mg/kg Tamoxifen (or corn oil only as a control) was injected intraperitoneally for 5 consecutive days, once a day.

### Balance beam experiments

Mice were trained on a balance beam two times a day for at least two days. The beam was 12 mm thick and had a dark box at the end with food and bedding. On the day of testing, the time to cross the beam was measured or the time and distance until the mouse fell was recorded. The number of times the paw slipped off the top of the beam was also recorded. For the tamoxifen induced model, a 6 mm thick beam was used.

### Footprints experiments

Mouse paws were pressed firmly into a blue ink pad. The mouse was placed on a narrow walkway with 1mm filter paper on the bottom and then allowed to walk from one end to another. Footprints were photographed, and representative images are shown.

### Astrocyte cultures

Cerebral cortices were dissected from pups (P1-P2) and meninges were removed. The tissue was dissociated mechanically, incubated in serum free Dulbecco’s Modified Eagle’s Medium (DMEM, Gibco) with trypsin and DNase1 for 30 minutes at 37°C, and centrifuged. The tissue was then titrated and filtered through a 100 μm filter to ensure a single cell suspension. After re-centrifuging, the cells were resuspended in DMEM supplemented with 10% fetal bovine serum, penicillin/streptomycin, and non-essential amino acids. The cells were plated onto dishes coated with poly-D-lysine in phosphate buffered saline (1 mg/ml), incubated at 37 °C with 5% CO_2_, and cultured for two weeks.

### Purification and culture of cerebellar neurons

Glia-free purified cerebellar neurons were isolated using immunopanning as described in (Risher, Patel et al., 2014). Briefly, cerebellar tissue was dissected out from P6 mouse brains and digested using papain-dissociation system from Worthington following manufacturer’s protocol (Worthington Biochem. Corp.). Triturated cells were applied to negative panning dishes to remove unwanted cells and cell debris. Cell suspension were incubated with positive panning dishes (coated with anti-L1, Millipore) followed by a gentle wash to remove un-attached non-neuronal cells. Pure cerebellar neurons were lifted from the plate and collected by centrifugation, cultured in serum free Neurobasal-SATO medium, containing BDNF, CNTF, and forskolin (Risher et al., 2014). After two days, half of the media was replaced with new NB-SATO medium containing AraC (4 µM). After 48 hours, media was replaced with NB-SATO (without AraC). Cells were cultured for 7 additional days.

### Western blotting

Cells or flash frozen tissue were lysed in 10 mM Tris (pH 7.4), 150 mM sodium chloride, 1 mM EDTA, 1% Nonidet P-40, 1% Triton X-100, 1 mM sodium orthovanadate, and Roche protease inhibitor mixture. Samples were separated on a 10% gel and transferred onto nitrocellulose membrane (GE Healthcare). Anti-β-tubulin (Santa Cruz Biotechnology; sc-9104), and anti-YY1 (Bethyl Laboratories; 779) primary antibodies were used. Antigen-antibody complexes were visualized by enhanced chemiluminescence using Immobilon Western blotting kit (Millipore).

### Quantitative PCR

Total RNA was prepared from flash frozen tissue with Trizol (Life Technologies), reversed transcribed with the high-capacity cDNA kit (Applied Biosystems), and amplified on the BioRad CFX Connect Real-time System. SYBR Green intron-spanning pre-designed qPCR primers (BioRad) were used. Gene expression levels were normalized to GAPDH and represented as fold change in expression relative to the control.

### Histology

Animals were perfused with 0.9% Sodium Chloride and then with 4% paraformaldehyde. Brains were post-fixed for 48 hours. Tissue was paraffin-embedded, sectioned (5 µm), and stained by H&E. Slides were imaged using the Vectra Polaris Imaging system at the Cancer Mouse Models Core Facility (VCU, Richmond, VA).

### Immunofluorescence

Animals were perfused with 0.9% Sodium Chloride and then with 4% paraformaldehyde. Brains were post-fixed overnight, and then cryopreserved with 30% sucrose in PBS for 48 hours at 4°C, embedded in optimal cutting medium (Tissue-Tek VWR), and 40 µm frozen sections were prepared on probe plus microscopy slides (Fisher Sci). Tissue was permeabilized in ice-cold acetone for 10 minutes, washed three times with 0.2% triton in PBS, and blocked in PBS with Triton (0.2%) and fish skin gelatin (Electron Microscopy). The primary antibodies, anti-GFAP (Cell Signaling; 1:300), anti-IBA (Wako Chemicals; 1:1000), anti-calbindin (Cell Signaling; 1:200), and anti-YY1 (Bethyl Laboratories; 779) were diluted in the blocking buffer and incubated overnight at 4°C. Subsequently, sections were washed three times, blocked, and incubated with Alexa Fluor 488 or Alexa Fluor 594 secondary antibodies (1:500, Invitrogen) for 90 minutes at room temperature. Nuclei were counterstained with Hoechst for 5 minutes at room temperature. Slides were washed three times and mounted using Vectashield mounting medium (Vector Laboratories). Slides were imaged using the Zeiss LSM 700 or the BZ-X800 (Keyence). Maximum projection images from z-stacks, and no fluorescence crossover was found between the channels. Images were analyzed using ImageJ. All locations were chosen for quantification when only viewing Hoechst. *GFAP^+^* processes were quantified in ImageJ by drawing lines along the entire length of the ML, the number of processes were counted, and the length of the line was measured. Purkinje cell processes were quantified by drawing a line across the PC layer. The length of the line was measured, and the number of PC bodies were counted. *IBA1^+^* cells were quantified in the entire area of ML, GL, or WM. The area was measured, and the number of cells were counted.

### Preparation of cerebellar slices

Brain slices were obtained by methods previously described (Bell, Shim et al., 2011). Mice were deeply anaesthetized with an intraperitoneal injection of ketamine (200 mg/kg) and xylazine (20 mg/kg), transcardially perfused with ice cold buffer consisting of 230 mM Sucrose, 3 mM KCl, 2 mM CaCl, 6 mM MgCl_2_, 1 mM NaHPO_4_, 25 mM NaHCO_3_, and 25 mM glucose. The brain was removed, cut on a parasagittal plane, and parasagittal slices containing the vermis of the cerebellum were cut at 350 mm on a Leica VT1200 (Leica Microsystems, Buffalo Grove, IL). Sections were incubated in a holding chamber maintained at 32°C. The holding chamber solution consisted of 125 mM NaCl, 3.0 mM KCl, 1.2 mM CaCl, 1.2 mM MgCl_2_, 1.2 mM NaHPO_4_, 25 mM NaHCO_3_, 25 mM glucose bubbled with 95% O_2_ / 5% CO_2_. Recordings were performed at 32-35°C.

### Electrophysiological measurements

Whole cell patch clamp recordings from cerebellar Purkinje cells were performed using patch pipettes pulled from borosilicate glass (8250 1.65/1.0 mm) on a Sutter P-1000 micropipette puller. Patch pipettes were filled with 140 mM KMeSO_4_, 8 mM NaCl, 2 mM MgATP, 0.1 mM NaGTP, and 10 mM HEPES. Membrane potentials were measured with a Model 2400 patch clamp amplifier (A-M Systems) and converted into a digital signal by a PCI-6040E A/D board (National Instruments). WCP Strathclyde Software was used to store and analyze membrane potential responses (courtesy of Dr. J Dempster, Strathclyde University, Glasgow, Scotland). The calculated junction potential (9.4 mV) was not compensated for in the analysis. Further analysis was performed using the Julia programming language (Julia Computing), OriginPro 2018 (OriginLab Corp.) and Graphpad Prism. Data were analyzed using WCP software and OriginPro 2018 for electrophysiological measurements. Statistics were performed using GraphPad Prism. Tests for normal distribution of data were performed using the Shapiro-Wilk test (*P*<0.05). Data that were normally distributed were tested by an ordinary one-way ANOVA with Bonferroni post hoc tests. Data that did not pass the Shapiro-Wilk test were analyzed using the Kruskal-Wallis tests. Differences were determined to be statistically significant for *P* values less than 0.05. All data was reported as the mean with error bars representing standard error of the mean (SEM).

### Isolation of single cells at P10 and P17

Cerebellum were isolated from mice at P10 or P17, and meninges were removed in cold ACSF (120 mM NaCl, 3 mM KCl, 2 mM MgCl, 0.2 mM CaCl, 26.2 mM NaHCO_3_, 11.1 mM glucose, 5 mM HEPE, 3 mM AP5, and 3 mM CNQX) bubbled with 95% oxygen. The tissue was cut 15 times and dissociated with the Worthington Papain Dissociation kit (# LK003153). The manufactures protocol was followed. Briefly, the tissue was dissociated in the papain DNase solution for 30 minutes at 37°C with 5% CO_2_, gently swirling every 5 minutes. The cells were filtered through a 70 µm filter (BD Falcon) to ensure a single cell suspension. At P17 cell debris was removed with the Debris Removal Solution (Miltenyi Biotec), following the manufacturer’s protocol. The cells were resuspended in cold PBS with 0.5% BSA, and the cell viability and concentration were determined using the TC20 Automated Cell Counter (Biorad). The cells were diluted to a final concentration of 1x10^6^ cells/ml.

### single-cell RNA sequencing

There was a targeted cell recovery of 5000 single cells per sample, which was loaded onto the Chromium Controller (10x Genomics). The libraries were made with the Chromium Single Cell 3’ Library and Gel Bead Kit (10x Genomics), following the manufacturers protocol. Unique Chromium i7 Sample Indexes (10xGenomics) were added to each sample, following the manufacturers protocol. Prior to sequencing, all four samples were combined. Samples were sequenced on the NextSeq 500 with the 150 cycle kit using the following mode: 28 bp (Read 1), 8 bp (Index), and 91 bp (Read 2). FASTQ files were generated with 10x Genomics CellRanger after trimming raw reads from 3’ end to recommended 28 base-pair (bp) long R1 reads and 91bp long R2 reads. At P10 sequencing depth averaged to 97,000,509 raw reads per sample, with almost 29,000 reads per cell on average. At P17 the sequencing depth averaged to 113,023,689 with 52,298 reads per cell on average.

### scRNA-seq data processing

scRNA-seq data was processed with 10x Genomics software CellRanger (v 3.0.2). The function count with the option --expect-cells set to 5000 was used on each sample individually. Mouse genome version mm10 was used as a reference (v. mm10-3.0.0, downloaded from https://support.10xgenomics.com/single-cell-gene-expression/software/downloads/latest). Following alignment, barcode assignment and unique molecular identifier (UMI) counting, samples were merged using aggr function with the option --normalize=mapped. Downstream analysis was performed with R package Seurat (Butler, Hoffman et al., 2018, Stuart, Butler et al., 2019) using as input pre-filtered CellRanger gene-cell matrix outs/filtered_feature_bc_matrix/. The initial UMI matrix contained 31,053 unique genes. There were 13,778 cells from all four samples at P10 and 8,722 cells from all four samples at P17.

### Data filtering

Genes with no expression in any cell were removed (9,365 in total in P10, and 9,153 in P17). To remove presumably dead cells, mitochondrial gene expression ratio was calculated, and cells with mitochondrial gene expression ratio greater than four median absolute deviations (MAD) from the median value were removed. Upon inspection of density plots, hard cut-offs were applied for removal of cells with respect to the number of expressed genes and number of sequencing reads. To account for empty droplets and multiplet cells at P10, cells with less than 500 and more than 20,000 UMIs were removed, respectively. At P17, cells with less than 500 and more than 50,000 UMIs were removed, respectively. Similarly, cells with less than 250 and more than 6,000 expressed genes were removed at P10. Cells with less than 250 and more than 8,000 expressed were removed at P17. The final P10 UMI matrix contained 21,197 expressed genes and 11,226 cells and was used in downstream analysis. The final P17 UMI matrix contained 21,659 expressed genes and 7,321 cells and was used in downstream analysis.

### Cell clustering and data visualization

Briefly, the seurat object was normalized with NormalizeData(). FindVariableFeatures() was used to identify 2,000 highly variable genes, followed by data scaling with ScaleData(), and principal component analysis (PCA) with RunPCA(). The cells were then clustered by applying FindNeighbors (seurat_norm, dims = 1:40) and FindClusters (seurat_norm, resolution = 0.4). To visualize the data, uniform manifold approximation and projection method, UMAP (Becht, McInnes et al., 2018), was first applied with the function RunUMAP() to perform dimensionality reduction using the computed first 40 principal components (PCs) and then DimPlot() was used for visualization of the results.

### Subclustering and gene expression analysis

At P10 Cluster 6 was selected for more detailed analysis. Briefly, seurat object was subsetted to contain only cells matching cluster 6 and clustering procedure was again applied on that cluster with the following commands:

seurat_norm.cluster.6 <- subset(seurat_norm, idents = 6)

seurat_norm.cluster.6 <- FindNeighbors(seurat_norm.cluster.6, dims = 1:40)

seurat_norm.cluster.6 <- FindClusters(seurat_norm.cluster.6, resolution = 0.4)

The above procedure identified 4 subclusters (subCluster = (Risher et al., 2014)). FindMarkers() was then applied to determine biomarkers of each subcluster with respect to cluster 6 only, i.e., FindMarkers(seurat_norm.cluster.6, ident.1 = subCluster, logfc.threshold = 0.25, test.use = “roc”, only.pos = TRUE). Differential gene expression (DE) analysis was performed between subclusters and between conditions using FindMarkers() with adequate specifications, e.g., FindMarkers(cluster6.0, ident.1 = “WT”, ident.2 = “cKO”, logfc.threshold = .50, test.use=“MAST”) for DE analysis of a subcluster between conditions and FindMarkers(seurat_norm.cluster.6, ident.1 = 0, ident.2 = 1, logfc.threshold = .50, test.use=“MAST”) for DE analysis between two subclusters. At P17 analogous analysis was performed for clusters 9 and 10.

### Venn Diagrams

At P10, the top significant genes of cluster 6 (compared to all other P10 clusters) were compared to previous mouse astrocyte data sets (Zhang, Sloan et al., 2016) (Batiuk, Martirosyan et al., 2020, Borggrewe, Grit et al., 2021). Overlapping DEGs were visualized in a Venn Diagrams, using the Intervene tool (Khan & Mathelier, 2017). Similarly, at P17 the top significant genes from each of the subgroups of astrocytes were compared to the published data sets indicated above.

### Metaneighbour analysis

An unsupervised version of Metaneighbour analysis was applied to astrocyte clusters to estimate similarities within and across the P10 and P17 subpopulations. The analysis was performed as outlined in “Analysis of single cell RNA-seq data” course, section 8.8.5 (http://hemberg-lab.github.io/scRNA.seq.course/). The script ’2017-08-28-runMN-US.R’ to perform the analysis was obtained from ’https://github.com/hemberg-lab/scRNA.seq.course/tree/master/course_files/utils’. In brief, P10 and P17 seurat objects were saved as SingleCellExperiment(). Selected P10 and P17 subclusters were then merged into a single expression matrix and most variable genes were identified. By running the function ’run_MetaNeighbor_US’, similarities of the clusters across the two datasets were calculated and visualized with heatmap().

### Data and codes availability

Raw sequencing data are publicly available through the GEO database (GSE166792). Analysis was performed using R Statistical Language and all the analysis codes are available upon request.

### Statistical analysis

Statistical analysis was performed using GraphPad Prism 9. Values are displayed as mean ± standard error. T tests or ANOVAs were performed as indicated.

## Supplementary files

**Supplementary videos 1-8**

**Videos 1 and 2:** *Yy1^loxP/loxP^* and *Yy1^ΔAST^* littermates in the cage.

**Videos 3-5:** *Yy1^loxP/loxP^* mice crossing the balance beam.

**Videos 6-8:** *Yy1^ΔAST^* mice crossing the balance beam.

---

**Supplementary Fig. 1.**
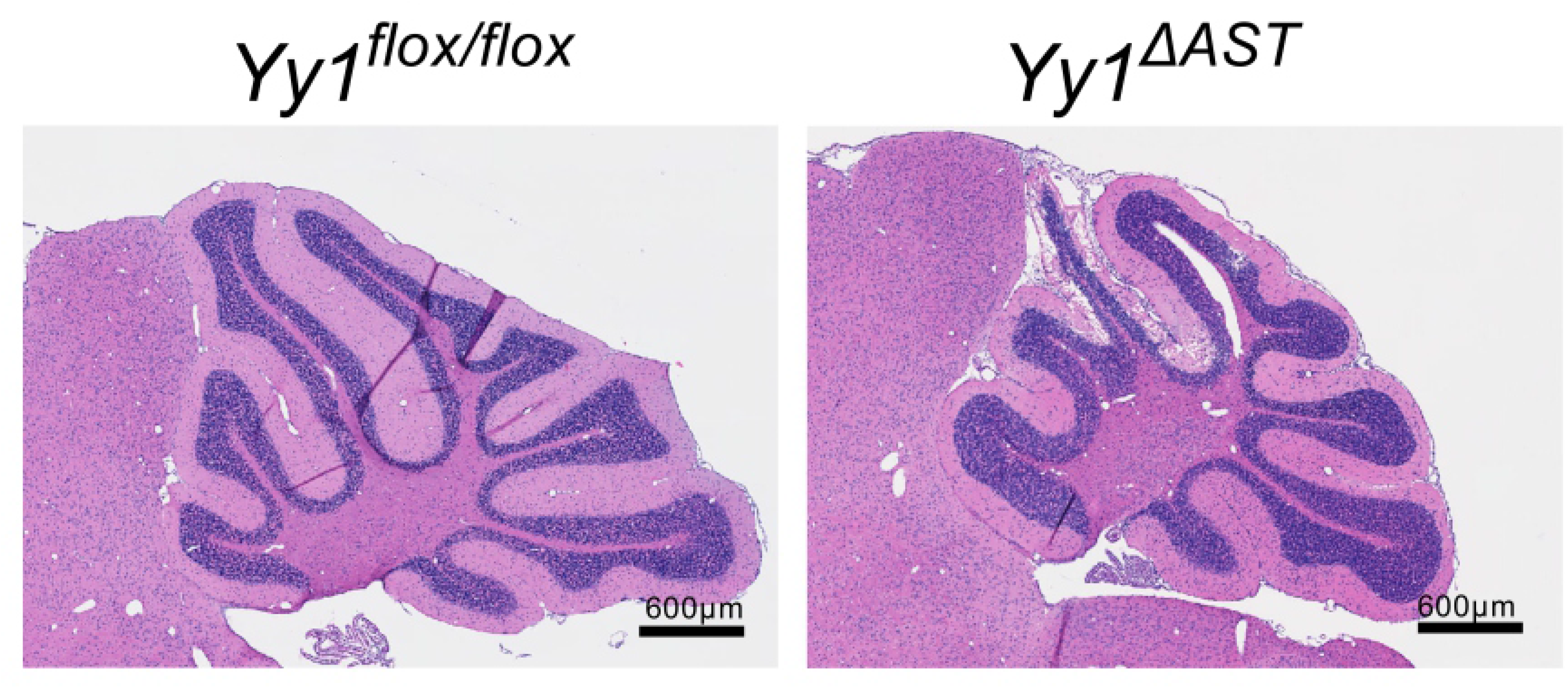
Cerebellar neurodegeneration in 5-month old *Yy1^ΔAST^* mice. Hematoxylin and Eosin staining of *Yy1^loxP/loxP^* and *Yy1^ΔAST^* 5-month old 8-12 cerebellum.

**Supplementary Fig. 2.**
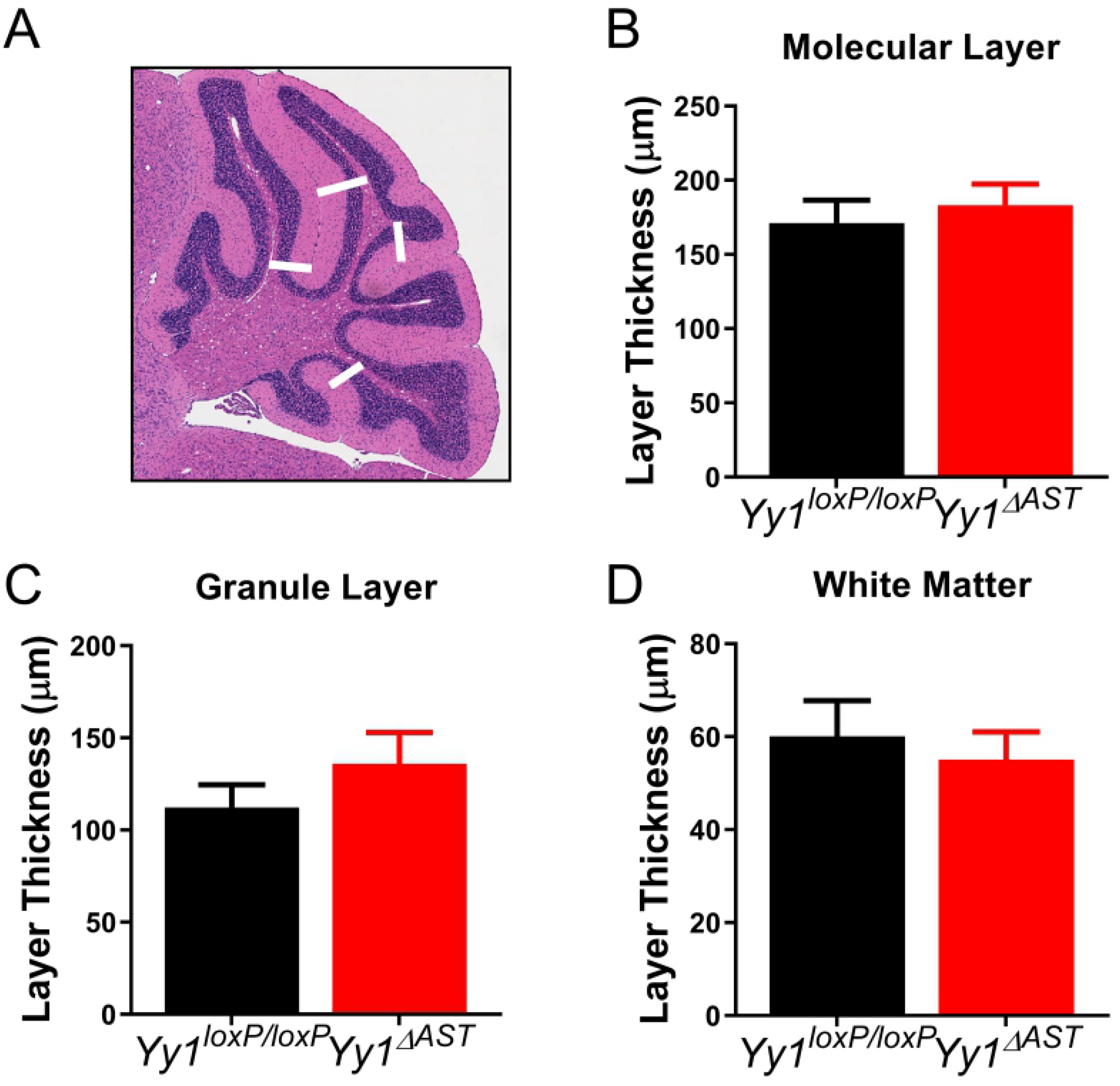
No difference in thickness of cerebellar layers in adult mice. (**A**) Locations measured in the cerebellum. (**B**) Mean ML thickness (n=12, 12 locations; two-tailed t-test, p=0.56). (**C**) Mean GL thickness (n=12, 12 locations; two-tailed t-test, p=0.27). (**D**) Mean WM thickness (n=12, 12 locations; two-tailed t-test, p=0.61).

**Supplementary Fig. 3.**
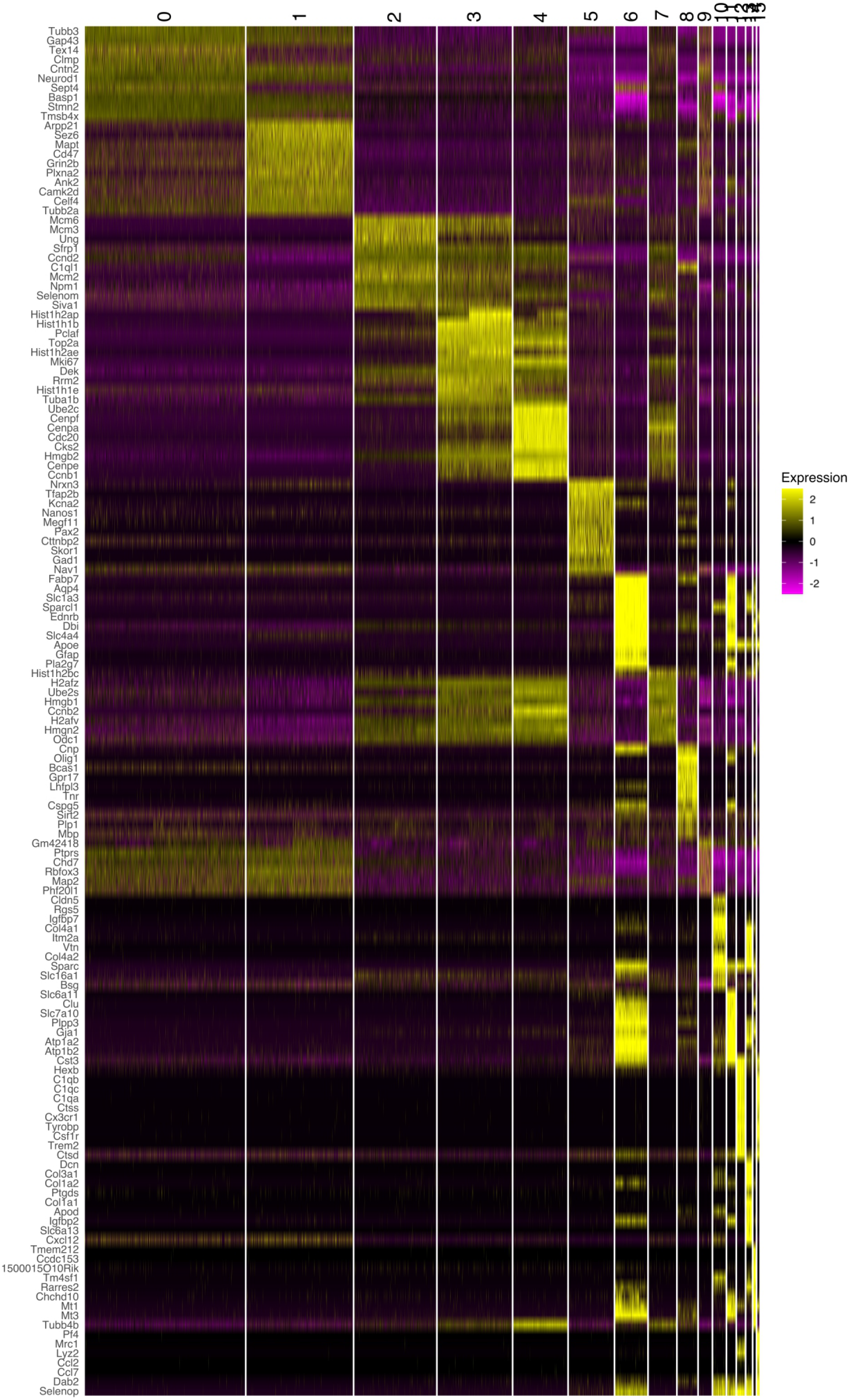
Heatmap of top 10 enriched genes at P10. Unsupervised analysis of gene expression profiles of the top 10 biomarkers for each cluster in *Yy1^LoxP/LoxP^* and *Yy1^ΔAST^* cerebella.

**Supplementary Fig. 4.**
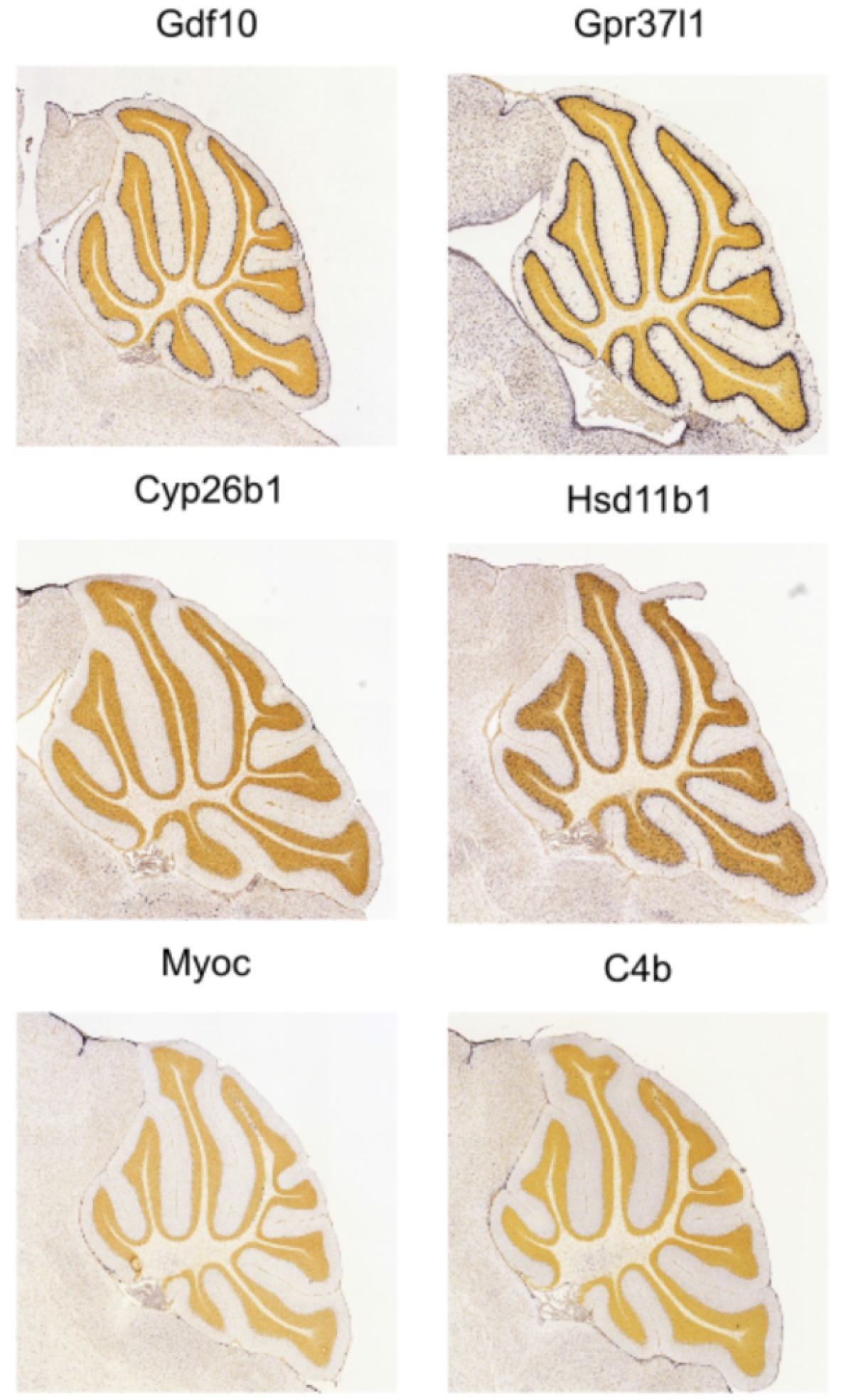
Allen brain atlas images at P28. *In situ* mRNA expression of identified subpopulation markers at P28.

**Supplementary Fig. 5.**
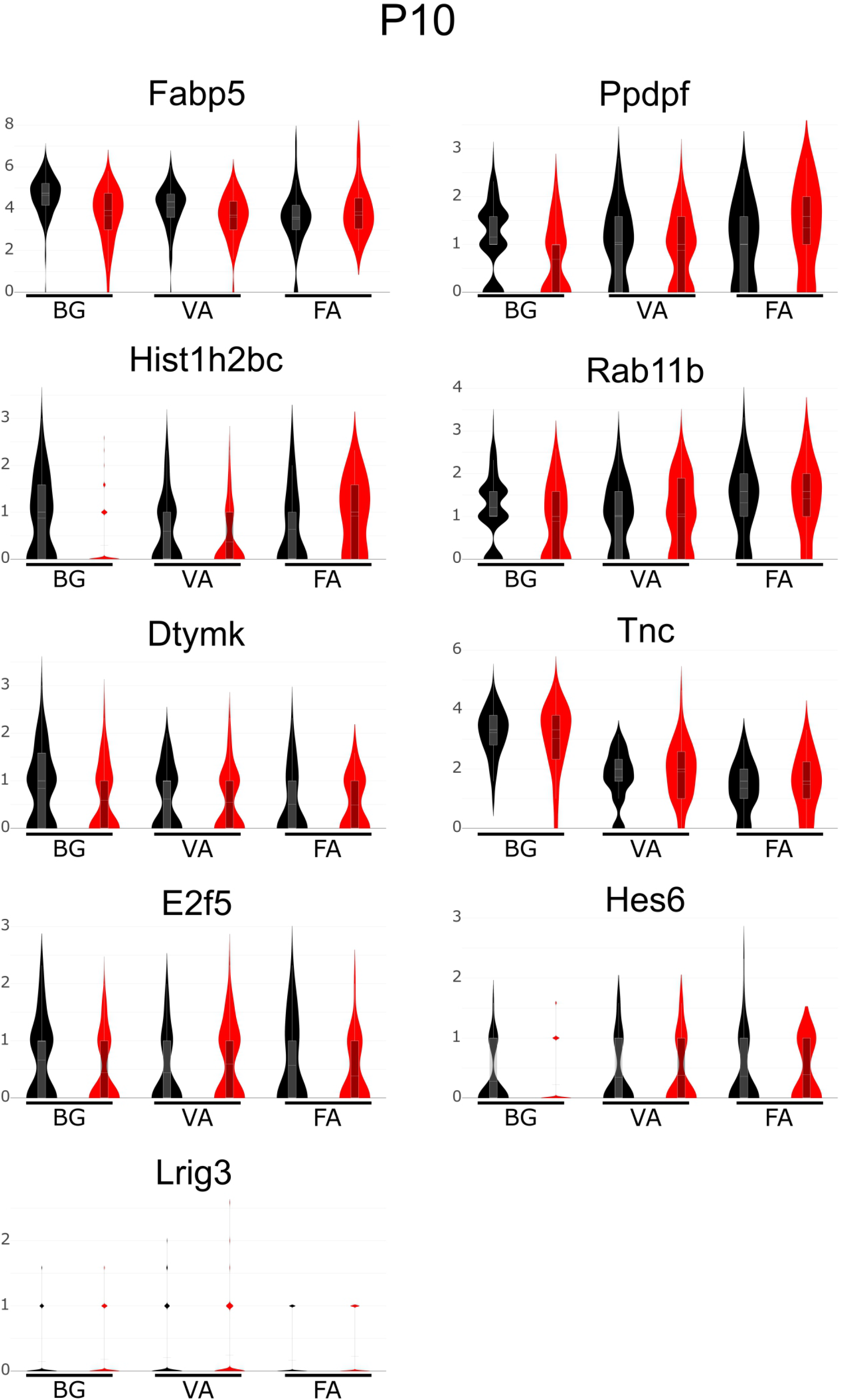
Expression of astrocyte progenitor markers by cerebellar astrocyte subpopulations at P10. Violin plot visualization of astrocyte progenitor marker expression (Zhang et al., 2016) in BG, VA, and FA at P10.

**Supplementary Fig. 6.**
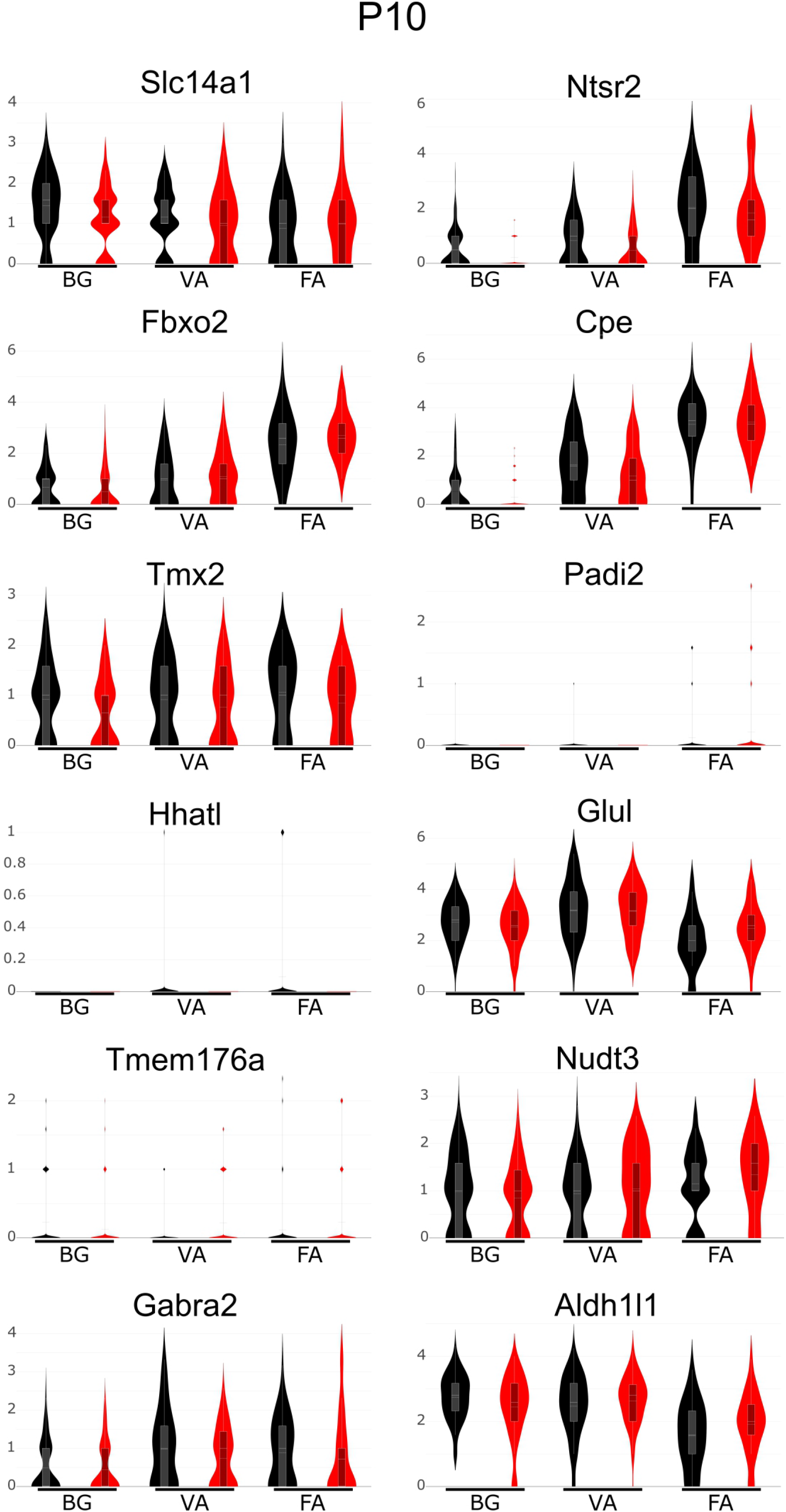
Expression of mature astrocyte markers by cerebellar astrocyte subpopulations at P10. Violin plot visualization of mature astrocyte marker expression (Zhang et al., 2016) in BG, VA, and FA at P10.

**Supplementary Fig. 7.**
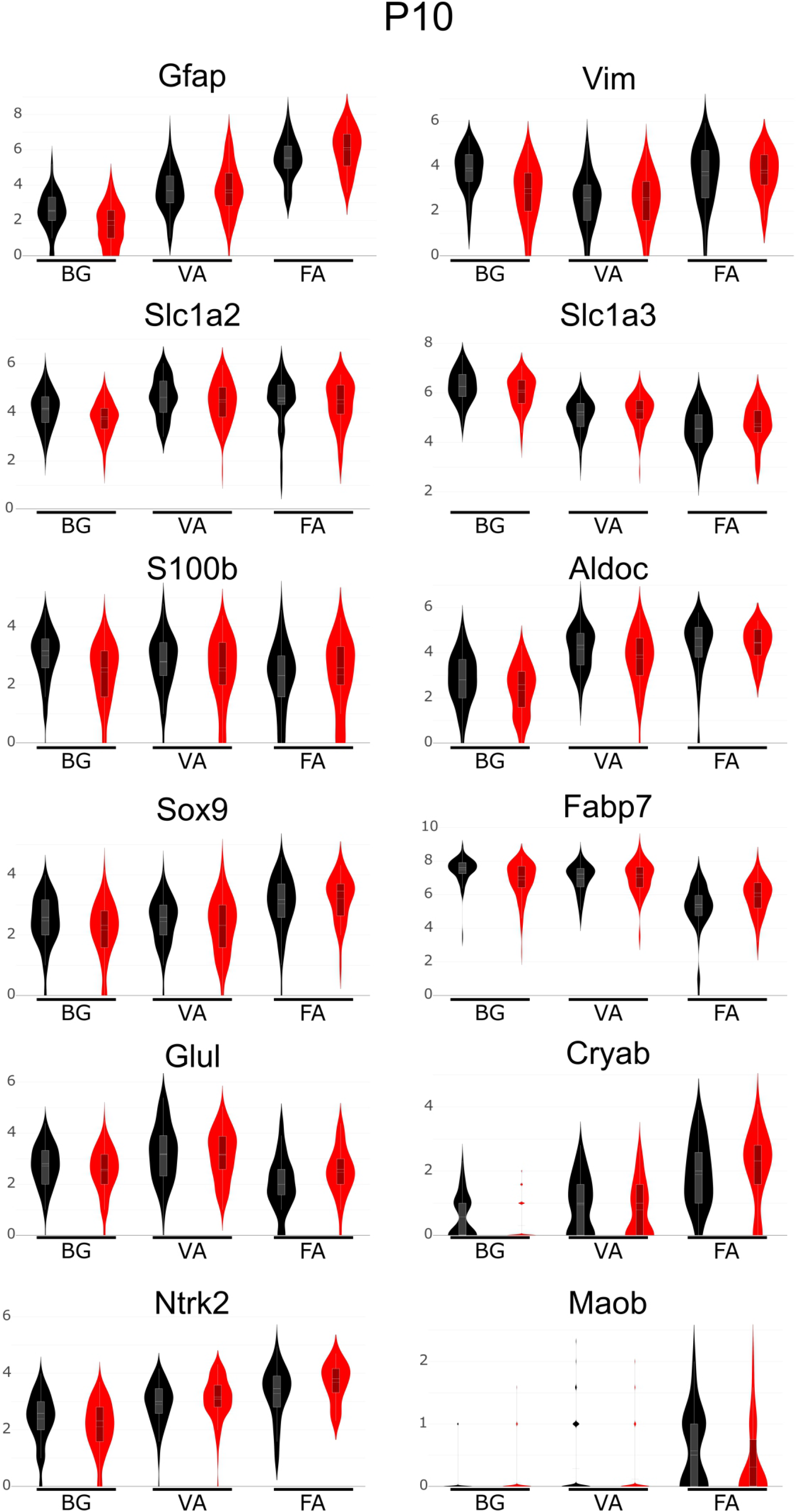
Expression of reactive astrocyte markers by cerebellar astrocyte subpopulations at P10. Violin plot visualization of reactive astrocyte marker expression (Escartin et al., 2021) in BG, VA, and FA at P10.

**Supplementary Fig. 8.**
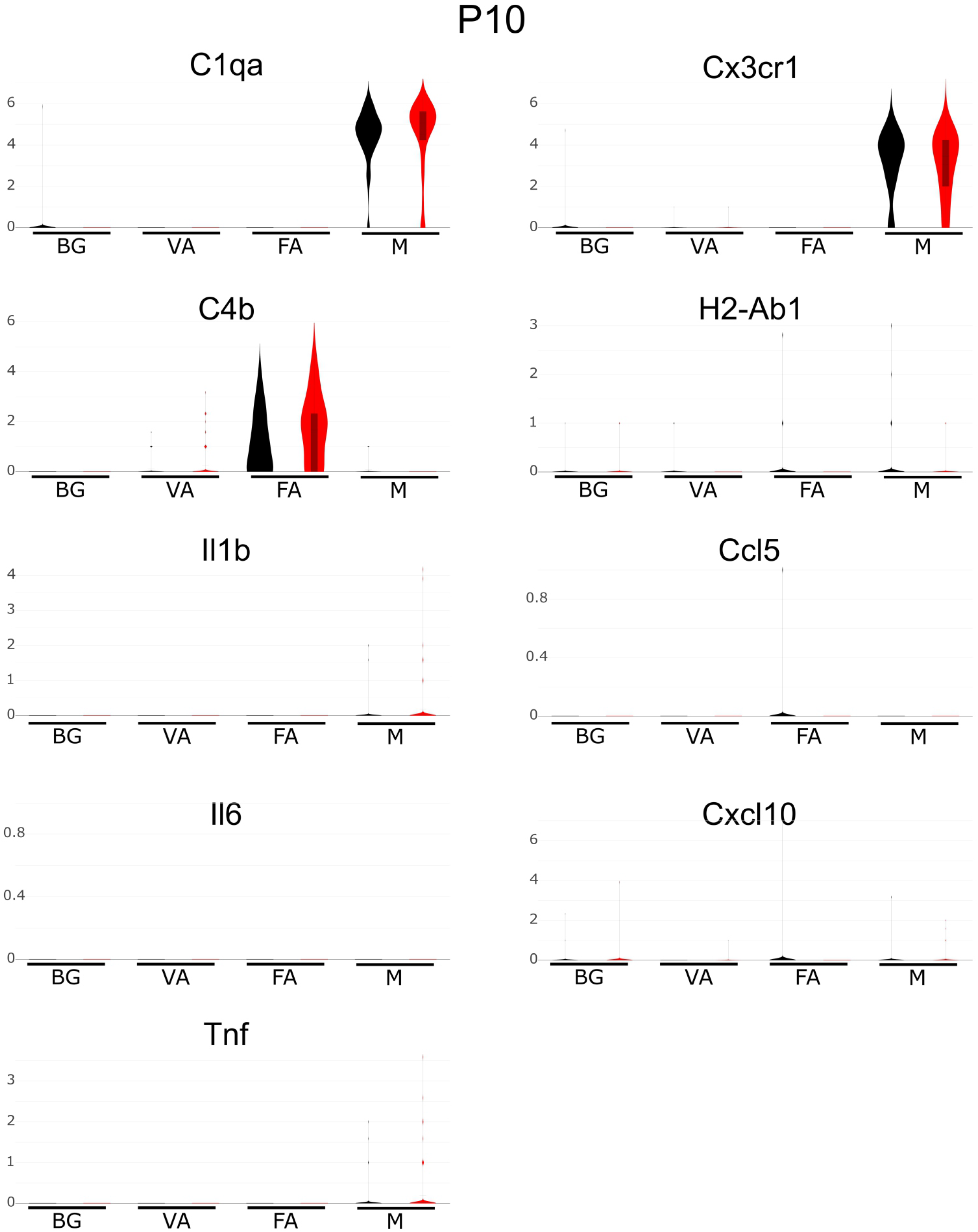
Expression of inflammatory mediators by astrocyte subpopulations and microglia in P10 cerebellum. Violin plot visualization of inflammatory mediators in BG, VA, FA, and microglia (M) at P10.

**Supplementary Fig. 9.**
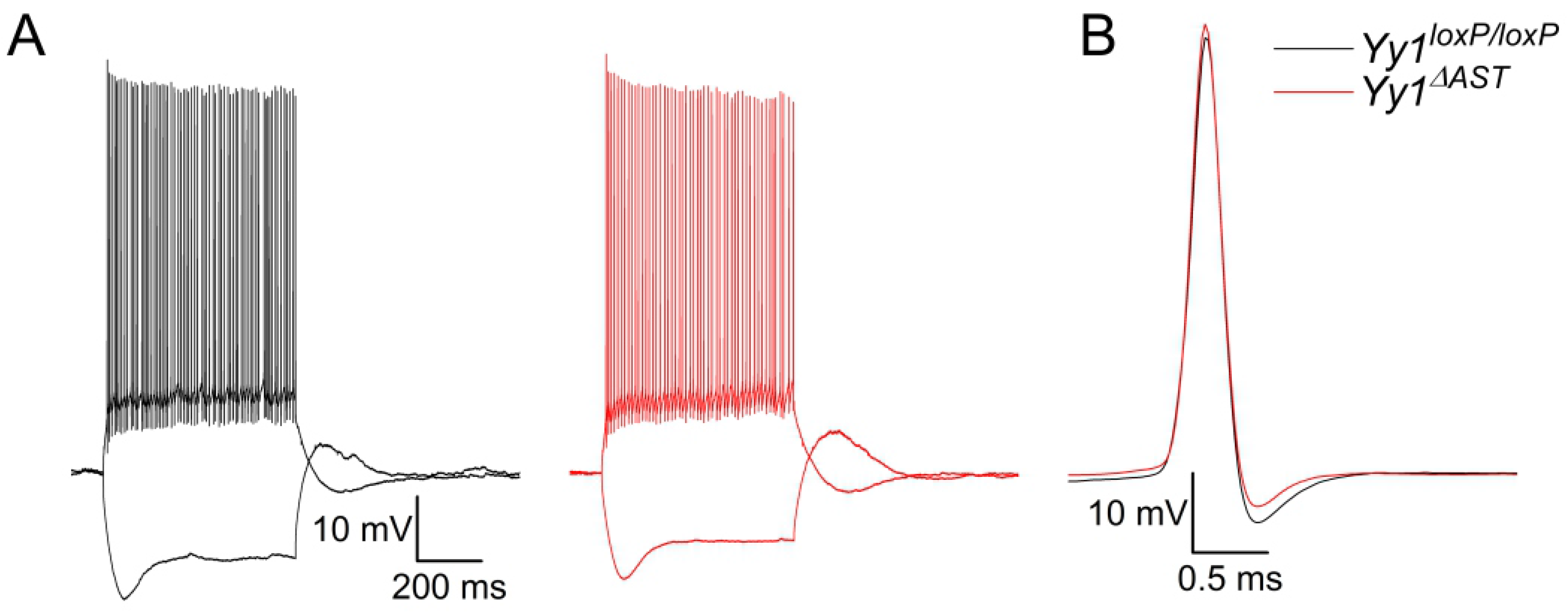
Electrophysiological properties of PC at P20. (**A**) Representative responses of a P20 *Yy1^loxP/loxP^* and *Yy1^ΔAST^* PC to 350 pA depolarizing and -400 pA hyperpolarizing step current injections (600 ms duration). (**B**) Overlay of the first action potentials in the train of A.

**Supplementary Fig. 10.**
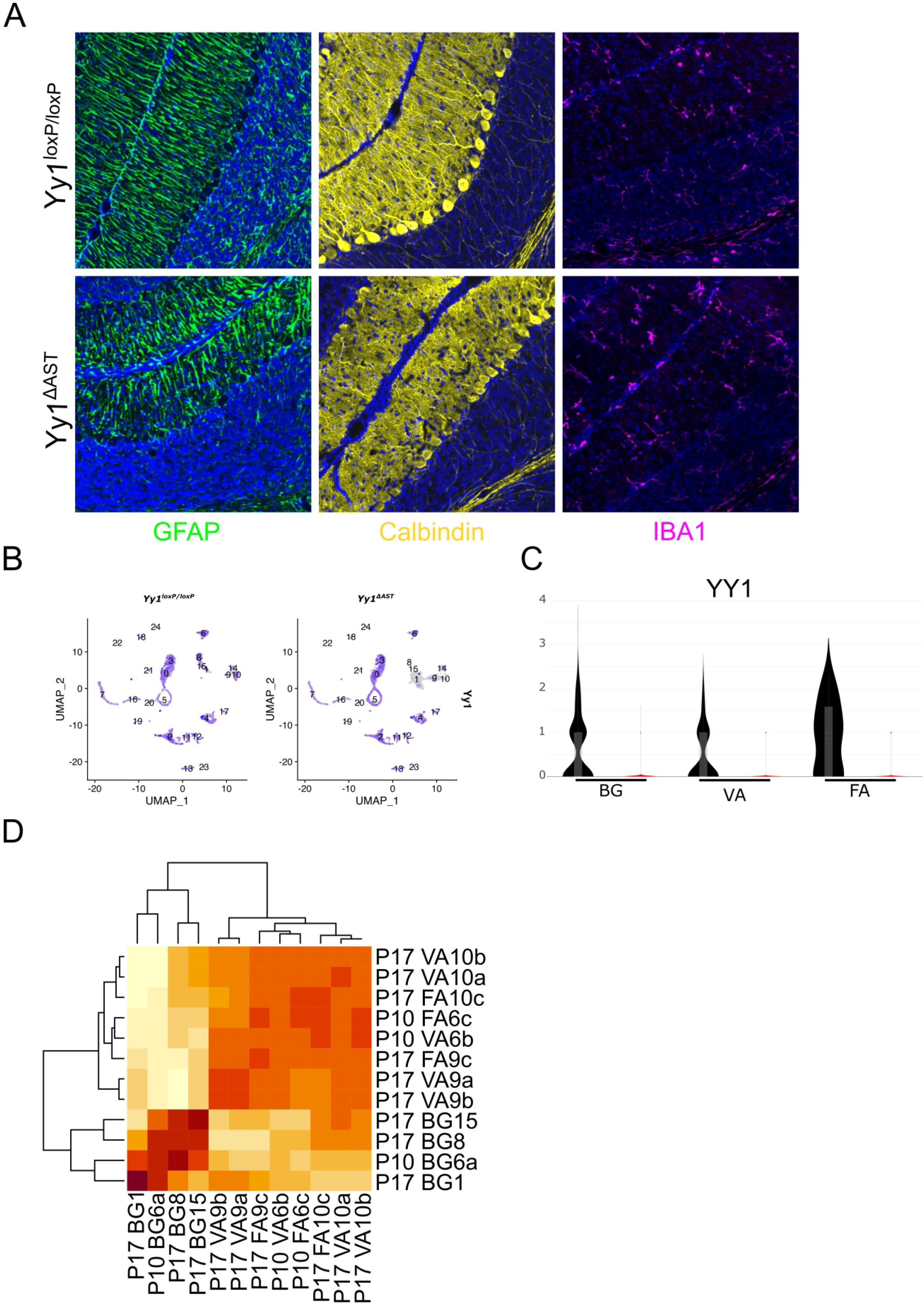
Divergent effects of YY1 on subpopulations of astrocytes in the developing cerebellum at P17. (**A**) Immunofluorescence of P17 cerebellum stained with GFAP, Calbindin, Iba1, and Hoechst; Images at 20X. ML, molecular layer; GCL, granular cell layer; WM, white matter. (**B**) UMAP and (**C**) violin plot visualization of YY1 expression in *Yy1^loxP/loxP^* and *Yy1^ΔAST^* cerebella. (**D**) Hierarchical clustering of P10 and P17 astrocyte clusters.

**Supplementary Fig. 11.**
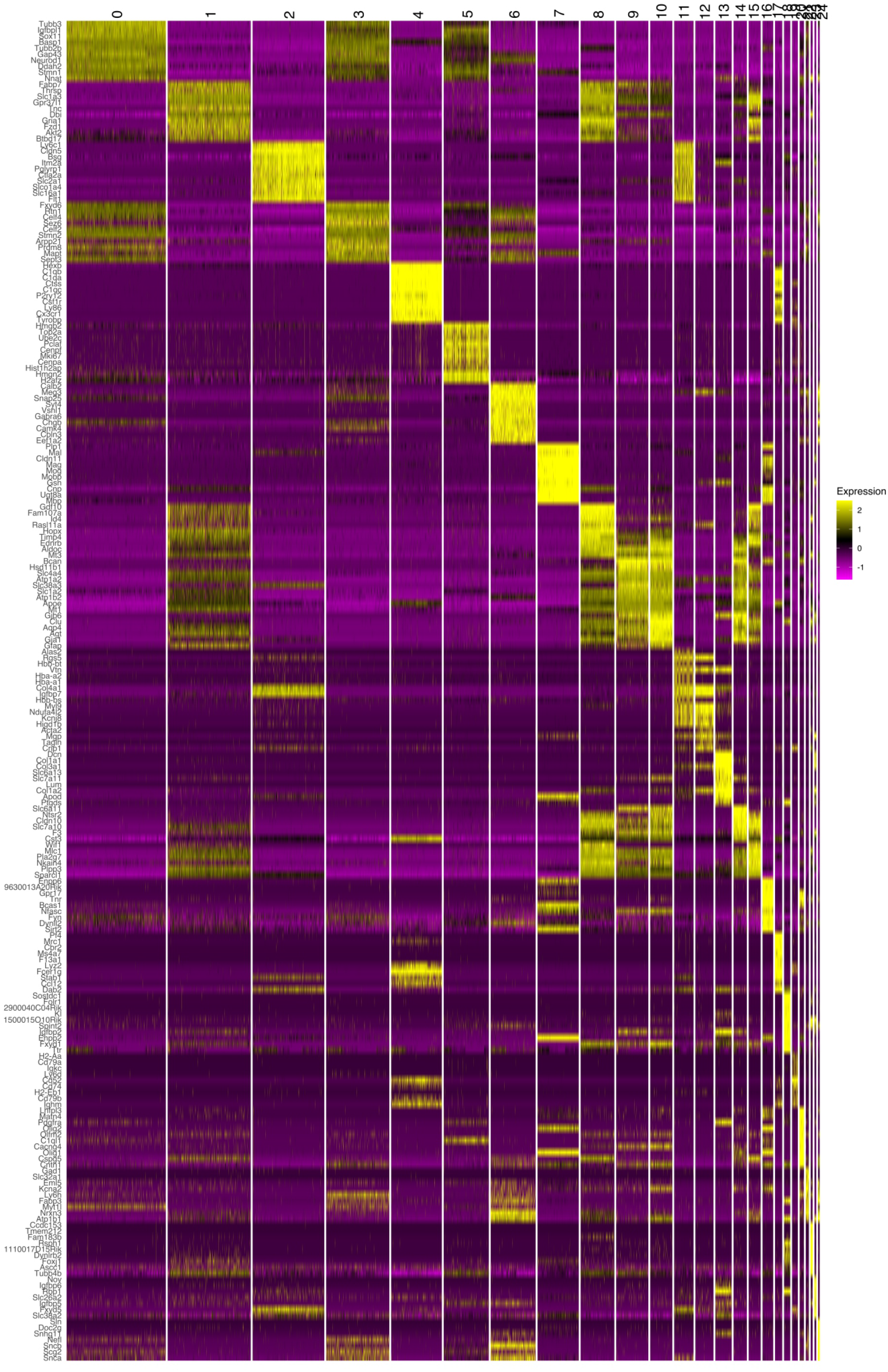
Heatmap of top 10 genes enriched at P17. Unsupervised analysis of gene expression profiles (top 10 biomarkers) for each cluster in *Yy1^LoxP/LoxP^* and *Yy1^ΔAST^* cerebella.

**Supplementary Fig. 12.**
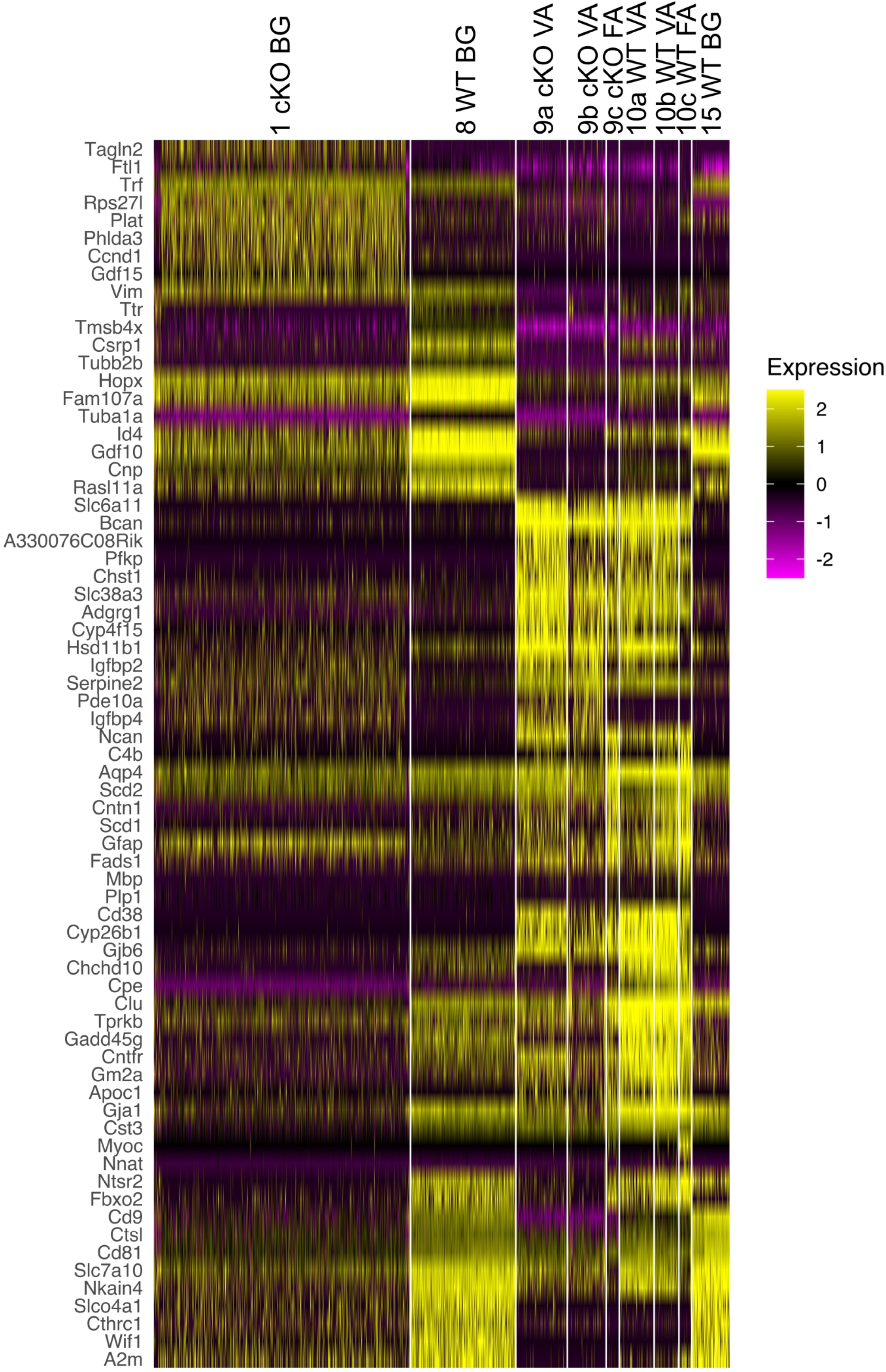
Heatmap of top 10 genes enriched between astrocyte clusters at P17. Heatmap (top 10 biomarkers for each subpopulation). Unsupervised analysis of gene expression profiles of BG, VA, and FA in *Yy1^LoxP/LoxP^* and *Yy1^ΔAST^* cerebella.

**Supplementary Fig. 13.**
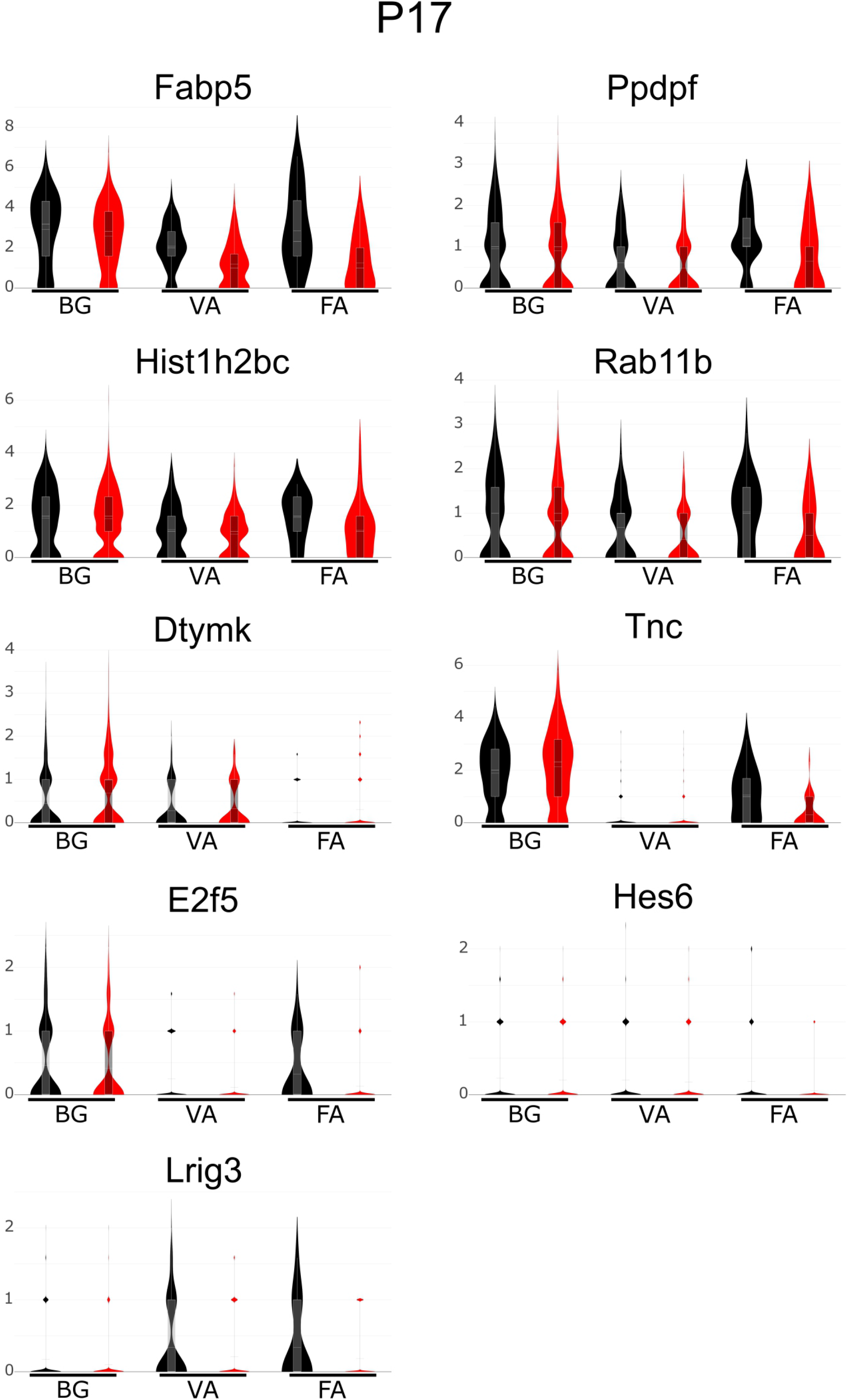
Expression of astrocyte progenitor markers by cerebellar astrocyte subpopulations at P17. Violin plot visualization of astrocyte progenitor marker expression (Zhang et al., 2016) in BG, VA, and FA at P17.

**Supplementary Fig. 14.**
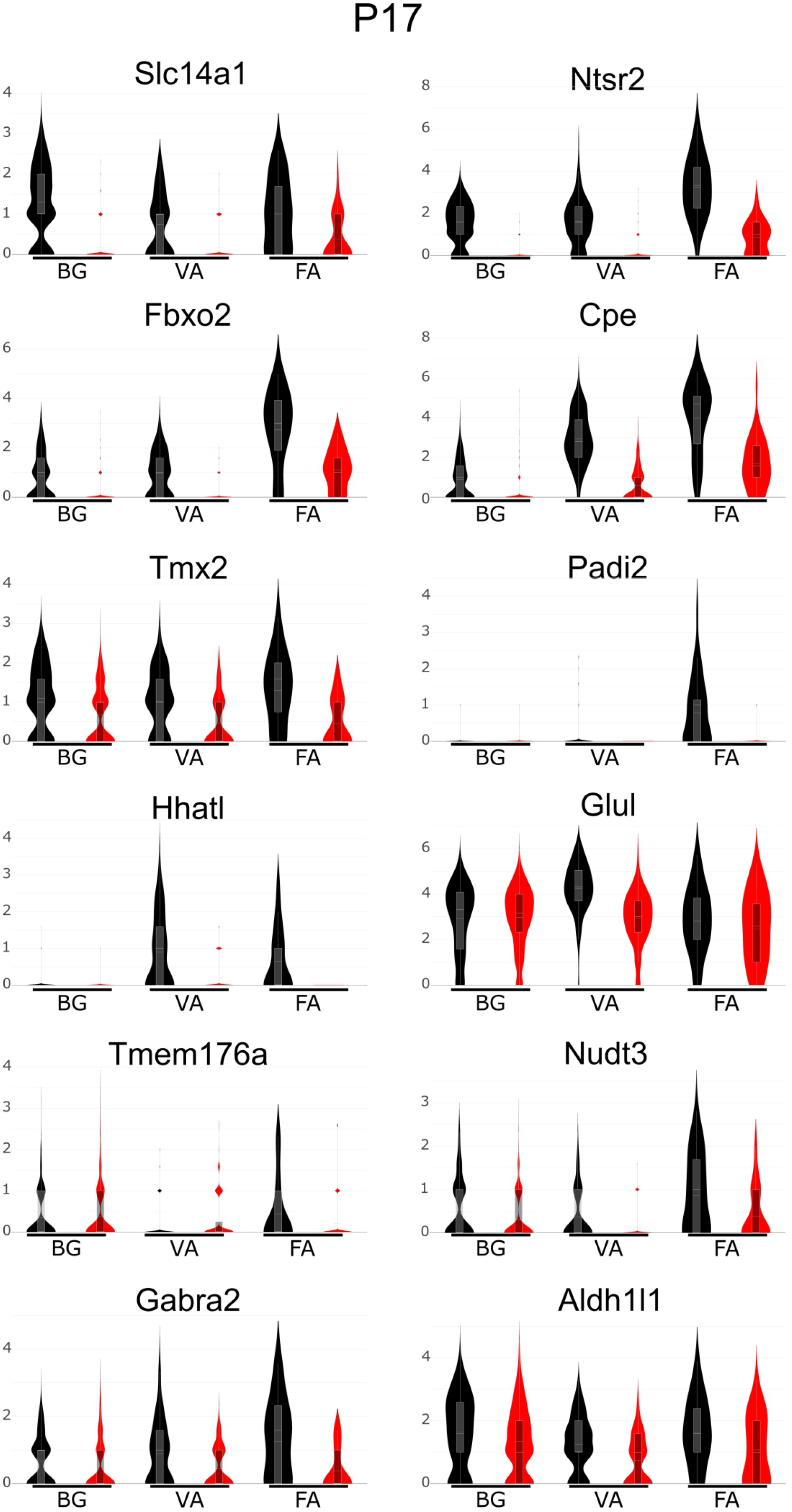
Expression of mature astrocyte markers by cerebellar astrocyte subpopulations at P17. Violin plot visualization of mature astrocyte marker expression (Zhang et al., 2016) in BG, VA, and FA at P17.

**Supplementary Fig. 15.**
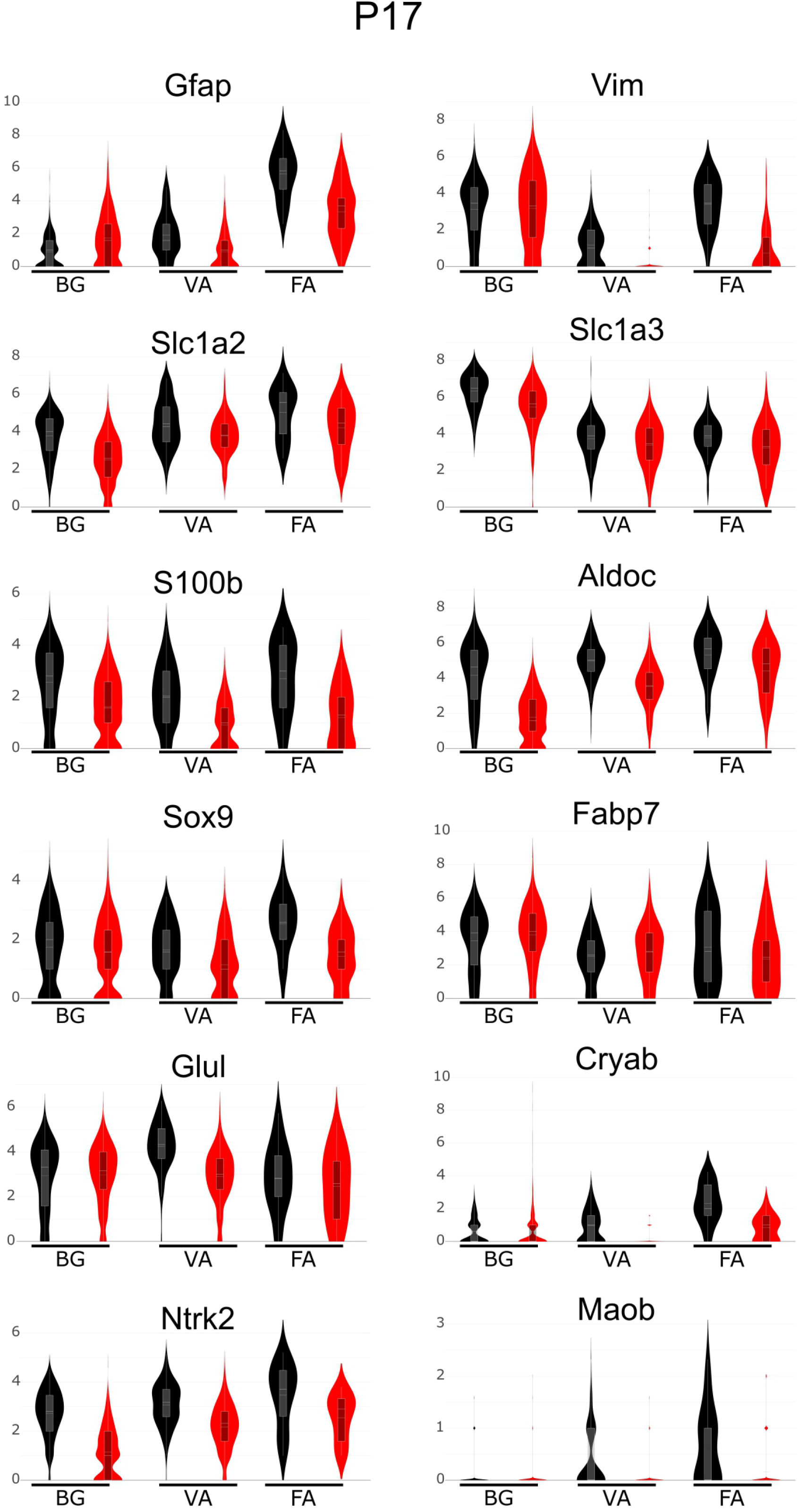
Expression of reactive astrocyte markers by cerebellar astrocyte subpopulations at P17. Violin plot visualization of reactive astrocyte marker expression (Escartin et al., 2021) in BG, VA, and FA at P17.

**Supplementary Fig. 16.**
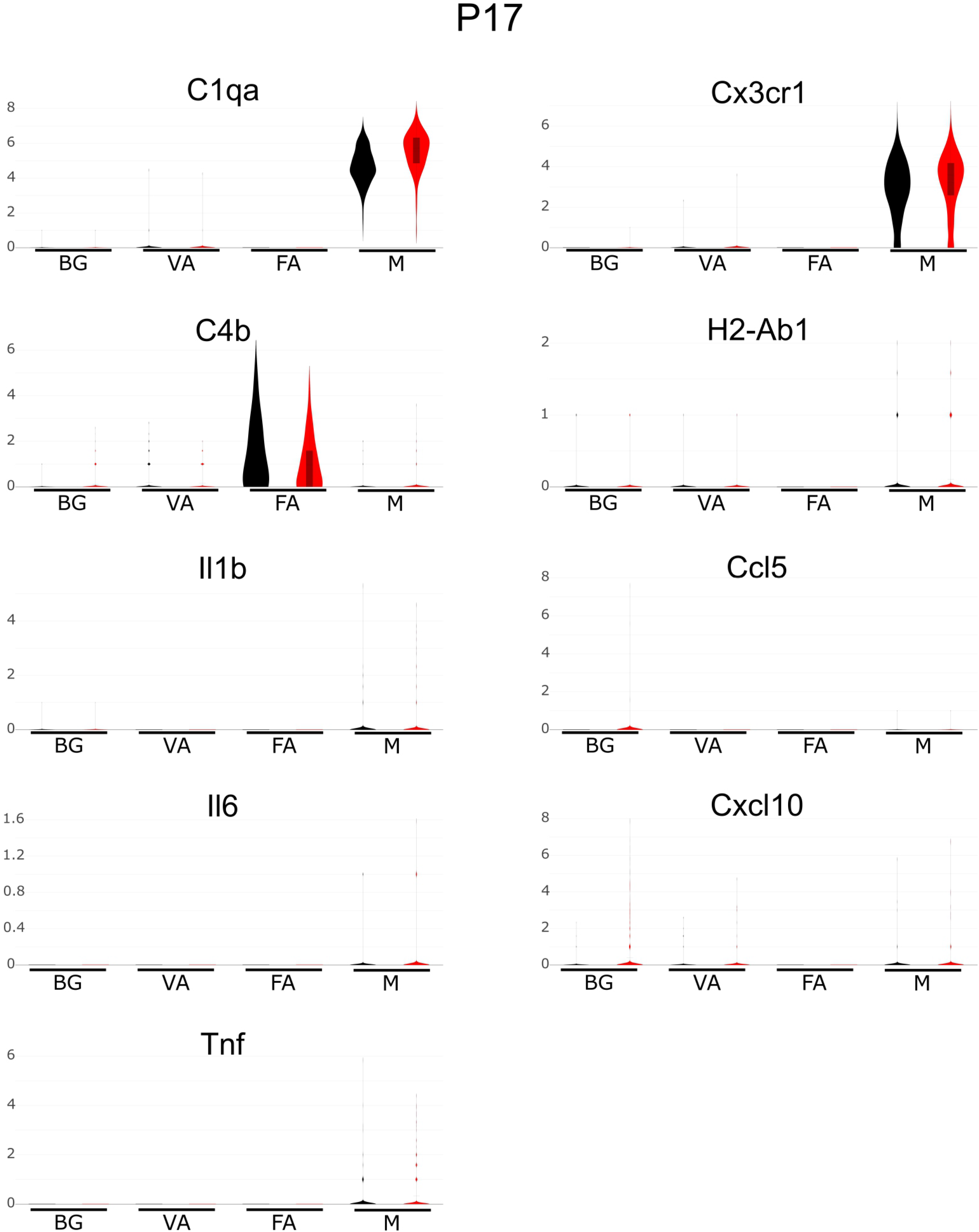
Expression of inflammatory mediators by astrocyte subpopulations and microglia in P17 cerebellum. Violin plot visualization of inflammatory mediators in BG, VA, FA, and microglia (M) at P17.

**Supplementary Table 1.** **Electrophysiological properties of *Yy1^ΔAST^* mice at P10.**

| Genotype | AP amplitude (mV) | AP duration (ms) | AP threshold (mV) | Mean Frequency (Hz) | Adaptation | Input Resistance (MΩ) | Sag Ratio |
| --- | --- | --- | --- | --- | --- | --- | --- |
| *Yy1<sup>LoxP/LoxP</sup>* | 35.9 ± 2.3 | 0.83 ± 0.05 | -36.8 ± 1.1 | 68.7 ± 8.1 | 0.80 ± 0.04 | 93.6 ± 6.9 | 0.53 ± 0.2 |
| *Yy1<sup>ΔAST</sup>* | 33.9 ± 2.7 | 0.77 ± 0.05 | -35.0 ± 1.3 | 51.9 ± 8.2 | 0.88 ± 0.06 | 106.1 ± 9.4 | 0.48 ± 0.01 |

**Supplementary Table 2.** **Summary of scRNA-seq at P10.** (**A**) The numbers of cells that distribute between different clusters and subclusters. (**B**) The top 10 biomarkers for each cluster as shown in Supplementary Fig. 3. (**C**) The top 10 biomarkers that distinguish *Yy1^LoxP/LoxP^* and *Yy1^ΔAST^* astrocyte clusters as shown in Fig. 4I. (**D**) The identified biomarkers between subgroups of astrocytes. (**E**) The identified DEGs between individual subgroups of astrocytes. (**F**) The DEGs between *Yy1^LoxP/LoxP^* and *Yy1^ΔAST^* astrocytes. (**G**) The DEGs between wildtype 6a versus both 6b and 6c. (**H**) The DEGs between wildtype 6b versus both 6a and 6c. (**I**) The DEGs between wildtype 6c versus both 6a and 6b.

**Supplementary Table 3.**
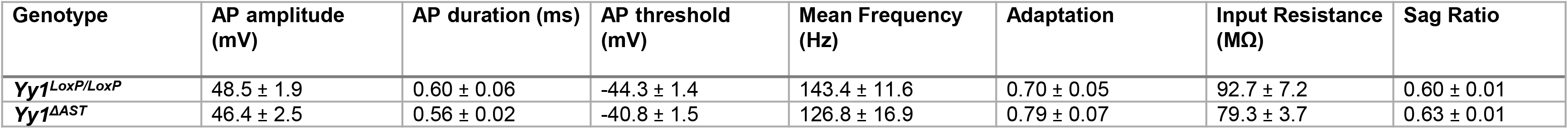
**Electrophysiological properties of *Yy1^ΔAST^* mice at P20.**

| Genotype | AP amplitude (mV) | AP duration (ms) | AP threshold (mV) | Mean Frequency (Hz) | Adaptation | Input Resistance (MΩ) | Sag Ratio |
| --- | --- | --- | --- | --- | --- | --- | --- |
| **Yy1**<sup>LoxP/LoxP</sup> | 48.5 ± 1.9 | 0.60 ± 0.06 | -44.3 ± 1.4 | 143.4 ± 11.6 | 0.70 ± 0.05 | 92.7 ± 7.2 | 0.60 ± 0.01 |
| **Yy1**<sup>ΔAST</sup> | 46.4 ± 2.5 | 0.56 ± 0.02 | -40.8 ± 1.5 | 126.8 ± 16.9 | 0.79 ± 0.07 | 79.3 ± 3.7 | 0.63 ± 0.01 |

**Supplementary Table 4.** **Summary of scRNA-seq at P17.** (**A**) The numbers of cells that distribute between different clusters and subclusters. (**B**) The top 10 biomarkers for each cluster as shown in Supplementary Fig. 7. (**C**) The top 10 biomarkers that distinguish *Yy1^LoxP/LoxP^* and *Yy1^ΔAST^* astrocyte clusters as shown in Supplementary Fig. 8. (**D**) The identified biomarkers between subgroups of astrocytes. (**E**) The identified DEGs between individual subgroups of astrocytes. (**F**) The DEGs between *Yy1^LoxP/LoxP^* and *Yy1^ΔAST^* astrocytes. (**G**) The DEGs between *Yy1^ΔAST^* cluster 1 versus *Yy1^LoxP/LoxP^* 8 and 15. (**H**) The DEGs between *Yy1^ΔAST^* cluster 9a and 9b versus *Yy1^LoxP/LoxP^* 10a and 10b. (**I**) The DEGs between *Yy1^ΔAST^* 9c versus *Yy1^LoxP/LoxP^* 10c.

## References

Araujo APB, Carpi-Santos R, Gomes FCA (2019) The Role of Astrocytes in the Development of the Cerebellum. Cerebellum 18: 1017–1035

Atchison ML (2014) Function of YY1 in Long-Distance DNA Interactions. Front Immunol 5: 45

Batiuk MY, Martirosyan A, Wahis J, de Vin F, Marneffe C, Kusserow C, Koeppen J, Viana JF, Oliveira JF, Voet T, Ponting CP, Belgard TG, Holt MG (2020) Identification of region-specific astrocyte subtypes at single cell resolution. Nat Commun 11: 1220

Beagan JA, Duong MT, Titus KR, Zhou L, Cao Z, Ma J, Lachanski CV, Gillis DR, Phillips-Cremins JE (2017) YY1 and CTCF orchestrate a 3D chromatin looping switch during early neural lineage commitment. Genome Res 27: 1139–1152

Ben Haim L, Rowitch DH (2017) Functional diversity of astrocytes in neural circuit regulation. Nat Rev Neurosci 18: 31–41

Bengani H, Mendiratta S, Maini J, Vasanthi D, Sultana H, Ghasemi M, Ahluwalia J, Ramachandran S, Mishra RK, Brahmachari V (2013) Identification and Validation of a Putative Polycomb Responsive Element in the Human Genome. PLoS One 8: e67217

Borggrewe M, Grit C, Vainchtein ID, Brouwer N, Wesseling EM, Laman JD, Eggen BJL, Kooistra SM, Boddeke E (2021) Regionally diverse astrocyte subtypes and their heterogeneous response to EAE. Glia 69: 1140–1154

Buffo A, Rossi F (2013) Origin, lineage and function of cerebellar glia. Prog Neurobiol 109: 42–63

Cai Y, Jin J, Yao T, Gottschalk AJ, Swanson SK, Wu S, Shi Y, Washburn MP, Florens L, Conaway RC, Conaway JW (2007) YY1 functions with INO80 to activate transcription. Nat Struct Mol Biol 14: 872–4

Carter RA, Bihannic L, Rosencrance C, Hadley JL, Tong Y, Phoenix TN, Natarajan S, Easton J, Northcott PA, Gawad C (2018) A Single-Cell Transcriptional Atlas of the Developing Murine Cerebellum. Curr Biol 28: 2910–2920 e2

Cerrato V (2020) Cerebellar Astrocytes: Much More Than Passive Bystanders In Ataxia Pathophysiology. J Clin Med 9

Cerrato V, Parmigiani E, Figueres-Onate M, Betizeau M, Aprato J, Nanavaty I, Berchialla P, Luzzati F, de’Sperati C, Lopez-Mascaraque L, Buffo A (2018) Multiple origins and modularity in the spatiotemporal emergence of cerebellar astrocyte heterogeneity. PLoS Biol 16: e2005513

Chai H, Diaz-Castro B, Shigetomi E, Monte E, Octeau JC, Yu X, Cohn W, Rajendran PS, Vondriska TM, Whitelegge JP, Coppola G, Khakh BS (2017) Neural Circuit-Specialized Astrocytes: Transcriptomic, Proteomic, Morphological, and Functional Evidence. Neuron 95: 531–549 e9

Chen CY, Shi W, Balaton BP, Matthews AM, Li Y, Arenillas DJ, Mathelier A, Itoh M, Kawaji H, Lassmann T, Hayashizaki Y, Carninci P, Forrest AR, Brown CJ, Wasserman WW (2016) YY1 binding association with sex-biased transcription revealed through X-linked transcript levels and allelic binding analyses. Sci Rep 6: 37324

Chiang CM, Roeder RG (1995) Cloning of an intrinsic human TFIID subunit that interacts with multiple transcriptional activators. Science 267: 531–6

Dong X, Kwan KM (2020) Yin Yang 1 is critical for mid-hindbrain neuroepithelium development and involved in cerebellar agenesis. Mol Brain 13: 104

Donohoe ME, Zhang X, McGinnis L, Biggers J, Li E, Shi Y (1999) Targeted disruption of mouse Yin Yang 1 transcription factor results in peri-implantation lethality. Mol Cell Biol 19: 7237–44

Escartin C, Galea E, Lakatos A, O’Callaghan JP, Petzold GC, Serrano-Pozo A, Steinhauser C, Volterra A, Carmignoto G, Agarwal A, Allen NJ, Araque A, Barbeito L, Barzilai A, Bergles DE, Bonvento G, Butt AM, Chen WT, Cohen-Salmon M, Cunningham C et al. (2021) Reactive astrocyte nomenclature, definitions, and future directions. Nat Neurosci 24: 312–325

Farmer WT, Abrahamsson T, Chierzi S, Lui C, Zaelzer C, Jones EV, Bally BP, Chen GG, Theroux JF, Peng J, Bourque CW, Charron F, Ernst C, Sjostrom PJ, Murai KK (2016) Neurons diversify astrocytes in the adult brain through sonic hedgehog signaling. Science 351: 849–54

Farmer WT, Murai K (2017) Resolving Astrocyte Heterogeneity in the CNS. Front Cell Neurosci 11: 300

Galloway A, Adeluyi A, O’Donovan B, Fisher ML, Rao CN, Critchfield P, Sajish M, Turner JR, Ortinski PI (2018) Dopamine Triggers CTCF-Dependent Morphological and Genomic Remodeling of Astrocytes. J Neurosci 38: 4846–4858

Ge WP, Miyawaki A, Gage FH, Jan YN, Jan LY (2012) Local generation of glia is a major astrocyte source in postnatal cortex. Nature 484: 376–80

Goertzen A, Veh RW (2018) Fananas cells-the forgotten cerebellar glia cell type: Immunocytochemistry reveals two potassium channel-related polypeptides, Kv2.2 and Calsenilin (KChIP3) as potential marker proteins. Glia 66: 2200–2208

Gomez CM (2019) The cerebellum in health and disease. Neurosci Lett 688: 1

Gordon S, Akopyan G, Garban H, Bonavida B (2006) Transcription factor YY1: structure, function, and therapeutic implications in cancer biology. Oncogene 25: 1125–42

Gregoire S, Karra R, Passer D, Deutsch MA, Krane M, Feistritzer R, Sturzu A, Domian I, Saga Y, Wu SM (2013) Essential and unexpected role of Yin Yang 1 to promote mesodermal cardiac differentiation. Circ Res 112: 900–10

Gupta I, Collier PG, Haase B, Mahfouz A, Joglekar A, Floyd T, Koopmans F, Barres B, Smit AB, Sloan SA, Luo W, Fedrigo O, Ross ME, Tilgner HU (2018) Single-cell isoform RNA sequencing characterizes isoforms in thousands of cerebellar cells. Nat Biotechnol

He L, Yu K, Lu F, Wang J, Wu LN, Zhao C, Li Q, Zhou X, Liu H, Mu D, Xin M, Qiu M, Lu QR (2018) Transcriptional Regulator ZEB2 Is Essential for Bergmann Glia Development. J Neurosci 38: 1575–1587

He Y, Dupree J, Wang J, Sandoval J, Li J, Liu H, Shi Y, Nave KA, Casaccia-Bonnefil P (2007) The transcription factor Yin Yang 1 is essential for oligodendrocyte progenitor differentiation. Neuron 55: 217–30

Hochstim C, Deneen B, Lukaszewicz A, Zhou Q, Anderson DJ (2008) Identification of positionally distinct astrocyte subtypes whose identities are specified by a homeodomain code. Cell 133: 510–22

Huang AY, Woo J, Sardar D, Lozzi B, Bosquez Huerta NA, Lin CJ, Felice D, Jain A, Paulucci-Holthauzen A, Deneen B (2020) Region-Specific Transcriptional Control of Astrocyte Function Oversees Local Circuit Activities. Neuron 106: 992–1008 e9

Ito K, Takizawa T (2018) Nuclear Architecture in the Nervous System: Development, Function, and Neurodevelopmental Diseases. Front Genet 9: 308

John Lin CC, Yu K, Hatcher A, Huang TW, Lee HK, Carlson J, Weston MC, Chen F, Zhang Y, Zhu W, Mohila CA, Ahmed N, Patel AJ, Arenkiel BR, Noebels JL, Creighton CJ, Deneen B (2017) Identification of diverse astrocyte populations and their malignant analogs. Nat Neurosci 20: 396–405

Karki P, Webb A, Smith K, Johnson J, Jr., Lee K, Son DS, Aschner M, Lee E (2014) Yin Yang 1 is a repressor of glutamate transporter EAAT2, and it mediates manganese-induced decrease of EAAT2 expression in astrocytes. Mol Cell Biol 34: 1280–9

Khakh BS (2019) Astrocyte-Neuron Interactions in the Striatum: Insights on Identity, Form, and Function. Trends Neurosci 42: 617–630

Khakh BS, Sofroniew MV (2015) Diversity of astrocyte functions and phenotypes in neural circuits. Nat Neurosci 18: 942–52

Kleiman E, Jia H, Loguercio S, Su AI, Feeney AJ (2016) YY1 plays an essential role at all stages of B-cell differentiation. Proc Natl Acad Sci U S A 113: E3911–20

Lee JS, Galvin KM, See RH, Eckner R, Livingston D, Moran E, Shi Y (1995) Relief of YY1 transcriptional repression by adenovirus E1A is mediated by E1A-associated protein p300. Genes Dev 9: 1188–98

Leto K, Arancillo M, Becker EB, Buffo A, Chiang C, Ding B, Dobyns WB, Dusart I, Haldipur P, Hatten ME, Hoshino M, Joyner AL, Kano M, Kilpatrick DL, Koibuchi N, Marino S, Martinez S, Millen KJ, Millner TO, Miyata T et al. (2016) Consensus Paper: Cerebellar Development. Cerebellum 15: 789–828

Liu H, Schmidt-Supprian M, Shi Y, Hobeika E, Barteneva N, Jumaa H, Pelanda R, Reth M, Skok J, Rajewsky K, Shi Y (2007) Yin Yang 1 is a critical regulator of B-cell development. Genes Dev 21: 1179–89

Lozzi B, Huang TW, Sardar D, Huang AY, Deneen B (2020) Regionally Distinct Astrocytes Display Unique Transcription Factor Profiles in the Adult Brain. Front Neurosci 14: 61

Marazziti D, Di Pietro C, Golini E, Mandillo S, La Sala G, Matteoni R, Tocchini-Valentini GP (2013) Precocious cerebellum development and improved motor functions in mice lacking the astrocyte cilium-, patched 1-associated Gpr37l1 receptor. Proc Natl Acad Sci U S A 110: 16486–91

Meyer LC, Paisley CE, Mohamed E, Bigbee JW, Kordula T, Richard H, Lutfy K, Sato-Bigbee C (2017) Novel role of the nociceptin system as a regulator of glutamate transporter expression in developing astrocytes. Glia 65: 2003–2023

Mo A, Mukamel EA, Davis FP, Luo C, Henry GL, Picard S, Urich MA, Nery JR, Sejnowski TJ, Lister R, Eddy SR, Ecker JR, Nathans J (2015) Epigenomic Signatures of Neuronal Diversity in the Mammalian Brain. Neuron 86: 1369–84

Morel L, Chiang MSR, Higashimori H, Shoneye T, Iyer LK, Yelick J, Tai A, Yang Y (2017) Molecular and Functional Properties of Regional Astrocytes in the Adult Brain. J Neurosci 37: 8706–8717

Neal M, Richardson JR (2018) Epigenetic regulation of astrocyte function in neuroinflammation and neurodegeneration. Biochim Biophys Acta Mol Basis Dis 1864: 432–443

Parmigiani E, Leto K, Rolando C, Figueres-Onate M, Lopez-Mascaraque L, Buffo A, Rossi F (2015) Heterogeneity and Bipotency of Astroglial-Like Cerebellar Progenitors along the Interneuron and Glial Lineages. J Neurosci 35: 7388–402

Pavlou MAS, Grandbarbe L, Buckley NJ, Niclou SP, Michelucci A (2019) Transcriptional and epigenetic mechanisms underlying astrocyte identity. Prog Neurobiol 174: 36–52

Petkova V, Romanowski MJ, Sulijoadikusumo I, Rohne D, Kang P, Shenk T, Usheva A (2001) Interaction between YY1 and the retinoblastoma protein. Regulation of cell cycle progression in differentiated cells. J Biol Chem 276: 7932–6

Ravi B, Kannan M (2013) Epigenetics in the nervous system: An overview of its essential role. Indian J Hum Genet 19: 384–91

Rezai-Zadeh N, Zhang X, Namour F, Fejer G, Wen YD, Yao YL, Gyory I, Wright K, Seto E (2003) Targeted recruitment of a histone H4-specific methyltransferase by the transcription factor YY1. Genes Dev 17: 1019–29

Rodriques SG, Stickels RR, Goeva A, Martin CA, Murray E, Vanderburg CR, Welch J, Chen LM, Chen F, Macosko EZ (2019) Slide-seq: A scalable technology for measuring genome-wide expression at high spatial resolution. Science 363: 1463–1467

Saab AS, Neumeyer A, Jahn HM, Cupido A, Simek AA, Boele HJ, Scheller A, Le Meur K, Gotz M, Monyer H, Sprengel R, Rubio ME, Deitmer JW, De Zeeuw CI, Kirchhoff F (2012) Bergmann glial AMPA receptors are required for fine motor coordination. Science 337: 749–53

Sathyanesan A, Kundu S, Abbah J, Gallo V (2018) Neonatal brain injury causes cerebellar learning deficits and Purkinje cell dysfunction. Nat Commun 9: 3235

Srinivasan L, Pan X, Atchison ML (2005) Transient requirements of YY1 expression for PcG transcriptional repression and phenotypic rescue. J Cell Biochem 96: 689–99

Srinivasan R, Lu TY, Chai H, Xu J, Huang BS, Golshani P, Coppola G, Khakh BS (2016) New Transgenic Mouse Lines for Selectively Targeting Astrocytes and Studying Calcium Signals in Astrocyte Processes In Situ and In Vivo. Neuron 92: 1181–1195

Su Y, Shin J, Zhong C, Wang S, Roychowdhury P, Lim J, Kim D, Ming GL, Song H (2017) Neuronal activity modifies the chromatin accessibility landscape in the adult brain. Nat Neurosci 20: 476–483

Sultan FA, Day JJ (2011) Epigenetic mechanisms in memory and synaptic function. Epigenomics 3: 157–81

Tao J, Wu H, Lin Q, Wei W, Lu XH, Cantle JP, Ao Y, Olsen RW, Yang XW, Mody I, Sofroniew MV, Sun YE (2011) Deletion of astroglial Dicer causes non-cell-autonomous neuronal dysfunction and degeneration. J Neurosci 31: 8306–19

Wang J, Wu X, Wei C, Huang X, Ma Q, Huang X, Faiola F, Guallar D, Fidalgo M, Huang T, Peng D, Chen L, Yu H, Li X, Sun J, Liu X, Cai X, Chen X, Wang L, Ren J et al. (2018) YY1 Positively Regulates Transcription by Targeting Promoters and Super-Enhancers through the BAF Complex in Embryonic Stem Cells. Stem Cell Reports 10: 1324–1339

Waters MR, Gupta AS, Mockenhaupt K, Brown LN, Biswas DD, Kordula T (2019) RelB acts as a molecular switch driving chronic inflammation in glioblastoma multiforme. Oncogenesis 8: 37

Weintraub AS, Li CH, Zamudio AV, Sigova AA, Hannett NM, Day DS, Abraham BJ, Cohen MA, Nabet B, Buckley DL, Guo YE, Hnisz D, Jaenisch R, Bradner JE, Gray NS, Young RA (2017) YY1 Is a Structural Regulator of Enhancer-Promoter Loops. Cell 171: 1573–1588 e28

Yamada H, Fredette B, Shitara K, Hagihara K, Miura R, Ranscht B, Stallcup WB, Yamaguchi Y (1997) The brain chondroitin sulfate proteoglycan brevican associates with astrocytes ensheathing cerebellar glomeruli and inhibits neurite outgrowth from granule neurons. J Neurosci 17: 7784–95

Zamanian JL, Xu L, Foo LC, Nouri N, Zhou L, Giffard RG, Barres BA (2012) Genomic analysis of reactive astrogliosis. J Neurosci 32: 6391–410

Zeisel A, Hochgerner H, Lonnerberg P, Johnsson A, Memic F, van der Zwan J, Haring M, Braun E, Borm LE, La Manno G, Codeluppi S, Furlan A, Lee K, Skene N, Harris KD, Hjerling-Leffler J, Arenas E, Ernfors P, Marklund U, Linnarsson S (2018) Molecular Architecture of the Mouse Nervous System. Cell 174: 999–1014 e22

Zhang Y, Barres BA (2010) Astrocyte heterogeneity: an underappreciated topic in neurobiology. Curr Opin Neurobiol 20: 588–94

Zhang Y, Sloan SA, Clarke LE, Caneda C, Plaza CA, Blumenthal PD, Vogel H, Steinberg GK, Edwards MS, Li G, Duncan JA, 3rd, Cheshier SH, Shuer LM, Chang EF, Grant GA, Gephart MG, Barres BA (2016) Purification and Characterization of Progenitor and Mature Human Astrocytes Reveals Transcriptional and Functional Differences with Mouse. Neuron 89: 37–53

Zurkirchen L, Varum S, Giger S, Klug A, Hausel J, Bossart R, Zemke M, Cantu C, Atak ZK, Zamboni N, Basler K, Sommer L (2019) Yin Yang 1 sustains biosynthetic demands during brain development in a stage-specific manner. Nat Commun 10: 2192

## References

Becht E, McInnes L, Healy J, Dutertre CA, Kwok IWH, Ng LG, Ginhoux F, Newell EW (2018) Dimensionality reduction for visualizing single-cell data using UMAP. Nat Biotechnol

Bell KA, Shim H, Chen CK, McQuiston AR (2011) Nicotinic excitatory postsynaptic potentials in hippocampal CA1 interneurons are predominantly mediated by nicotinic receptors that contain alpha4 and beta2 subunits. Neuropharmacology 61: 1379–88

Butler A, Hoffman P, Smibert P, Papalexi E, Satija R (2018) Integrating single-cell transcriptomic data across different conditions, technologies, and species. Nat Biotechnol 36: 411–420

Khan A, Mathelier A (2017) Intervene: a tool for intersection and visualization of multiple gene or genomic region sets. BMC Bioinformatics 18: 287

Risher WC, Patel S, Kim IH, Uezu A, Bhagat S, Wilton DK, Pilaz LJ, Singh Alvarado J, Calhan OY, Silver DL, Stevens B, Calakos N, Soderling SH, Eroglu C (2014) Astrocytes refine cortical connectivity at dendritic spines. Elife 3

Stuart T, Butler A, Hoffman P, Hafemeister C, Papalexi E, Mauck WM, 3rd, Hao Y, Stoeckius M, Smibert P, Satija R (2019) Comprehensive Integration of Single-Cell Data. Cell 177: 1888–1902 e21

